# Loss of presenilin 2 function age-dependently increases susceptibility to kainate-induced acute seizures and blunts hippocampal kainate-type glutamate receptor expression

**DOI:** 10.1101/2025.09.09.675240

**Authors:** Larissa Robinson-Cooper, Stephanie Davidson, Rami Koutoubi, Kangni Zhang, Hannah Park, Melissa Barker-Haliski

**Author notes:** co-first authors (Larissa Robinson-Cooper, Stephanie Davidson). Author contributions: CRediT LRC – Conceptualization; formal analysis; investigation; methodology; Writing – original draft; Writing – review and editing; SD – Conceptualization; formal analysis; investigation; methodology; Writing – original draft; Writing – review and editing; RK – investigation; Writing – review and editing; KZ – investigation; Writing – review and editing; HP - investigation; Writing – review and editing; MBH - Conceptualization; formal analysis; Funding acquisition; investigation; methodology; Project administration; Supervision; Writing – original draft; Writing – review and editing.

## Abstract

Presenilin 2 (PSEN2) variants increase risk of Alzheimer’s disease (AD) and unprovoked seizures. Yet, age-related PSEN2 contributions to seizure susceptibility are understudied. Critically, PSEN proteolytic capacity may regulate hippocampal kainate-type glutamate receptor (KARs) availability. Kainic acid (KA) is a KAR agonist that evokes severe seizures in mice. We hypothesized that PSEN2 knockout (KO) mice would show reduced latency to KA-induced seizures, increased seizure burden, worsened 7-day survival, and altered hippocampal KAR expression compared to wild-type (WT) controls. Using repeated low-dose systemic KA administration to 3–4- and 12–15-month-old PSEN2 KO versus WT mice, we quantified acute seizure latency and neuropathology. GluK2 and GluK5 KAR subunit expression was colocalized in astrocytes 7 days after seizures or sham to assess the impact of PSEN2 loss and seizures on hippocampal KARs. Young PSEN2 KO mice were more seizure-prone than WTs, while genotype did not change latency to first seizure. Aged females seized faster than young females and experienced greater mortality, unlike males. There was no difference in KAR subunit expression between young mouse genotypes and regardless of seizure history. In both genotypes, hippocampal CA3 astrocytes expressed GluK5 after seizures, however, astrocytic GluK2 upregulation only occurred in WTs. GluK5 expression was significantly reduced in aged seizure-naïve PSEN2 KO versus WT mice, while total GluK2 expression did not differ. Seizure-induced astrocytic GluK5 expression only occurred in WT mice in CA3, while astrocytic GluK2 expression occurred in both. Thus, PSEN2 loss may impair age-related hippocampal KAR expression, implicating KARs as understudied contributors to AD-related seizures.

**Highlights.:**

- Alzheimer’s disease (AD) and epilepsy share many pathophysiological links, but the contributions of distinct AD-risk factors are understudied.
- Loss of normal presenilin 2 function is associated with AD; its role in late-life seizures and neuropathology have been minimally evaluated.
- Loss of presenilin 2 increases susceptibility to kainic-acid induced acute seizures in young adult mice, mirroring late life risk.
- Increased susceptibility to kainic acid induced acute seizures due to loss of PSEN2 is lost in late life, suggesting earlier senescence.
- Loss of presenilin 2 function in aged mice is associated with reduced GluK5 expression and greater astrogliosis under basal conditions.

## 1. Introduction

Older adults with Alzheimer’s disease (AD) have a high risk of comorbid seizures. Seizure incidence is particularly high in those with early-onset AD (EOAD) and these seizures worsen functional decline^1–3^. People over age 65 represent the fastest growing demographic with epilepsy diagnosis, underscoring an urgent need to understand the conserved pathological mechanisms driving late life seizure risk. AD has long been hypothesized to be associated with pathological Aβ accumulation^4^; accordingly, most investigations have focused on the contributions of Aβ overproduction to disease progression. Several risk genes are implicated in EOAD, but variants in Amyloid Precursor Protein (APP), Presenilin 1 (PSEN1), and Presenilin 2 (PSEN2) are the most well-known^5^. PSEN1 or PSEN2 form the catalytic core of the γ-secretase complex, and APP is their most well-studied substrate^6^. Interestingly, APP duplication and PSEN1 variant models demonstrate increased seizure occurrence^7–9^, while the impacts of PSEN2 variants on seizure risk remain largely underexplored. This deficit is particularly concerning because these AD-associated genetic risk factors all equally influence clinical seizure risk^5^. Further, clinical PSEN1 variants are associated with AD onset as early as 39 years old, while PSEN2 variants tend to have later AD onset^10^. PSEN2 variants may even be masked in the general late-onset AD population^11^, potentially representing an understudied contributor to late-onset AD. As a result of this narrow focus on APP and PSEN1, hyperexcitability studies have been largely conducted in APP-overexpressing mouse AD models (e.g., 5xFAD and APP/PS1)^7–9^. These mice exhibit rapid Aβ deposition, therefore, studies are primarily conducted in younger mice, limiting insight on how advanced age additively affects seizure risk in AD. PSEN2 null mice do not accumulate Aβ (**Supplemental Figure 1** and ^12^) , but do exhibit heightened neuroinflammation^13; 14^, offering an opportunity to understand the Aβ-independent contributions to seizure susceptibility in both AD and aging. Yet, few studies have investigated the seizure-specific impact of loss of normal PSEN2 function in aged rodents. We thus aimed to fill this knowledge gap by assessing how Aβ-independent factors drive disease progression to better inform late-onset AD treatment.

PSEN2 is a particularly intriguing, understudied driver of AD pathology and neuronal hyperexcitability. PSEN2 is the predominant γ-secretase in microglia and AD-associated variants in PSEN2 are associated with exaggerated microglial reactivity and a primed inflammatory phenotype in glial cells^13; 15; 16^. As inflammation is central to seizures in epilepsy ^17^, this phenotype may similarly influence seizure risk in AD. PSEN2 variant-derived astrocytes present with increased basal GFAP expression, mirroring normal aging^16; 18^. Taken together, this suggests that loss of normal PSEN2 function may accelerate the onset of reactive gliosis. Human PSEN2 variants lead to a loss of normal γ-secretase function, making PSEN2 knockout (KO) mice useful to initially assess how loss of normal PSEN2 function influences seizure risk. PSEN2 KO mice have favorable breeding and longevity^12,19^, providing an optimal platform on which to assess the influences of global PSEN2 deletion in an efficient, long-term manner. We have earlier reported age-related shift in sensitivity of PSEN2 KO mice to corneal kindling, a model of acquired chronic seizures and epileptogenesis^20^. In contrast, young PSEN2 KO mice have decreased electrically-evoked minimal clonic threshold, a model of acute limbic seizures, versus age-matched WT mice. Notably, PSEN2 KO mice also exhibit high-frequency oscillations, a biomarker of epilepsy, matching the occurrence in two other amyloidogenic AD models^21^. Thus, loss of normal PSEN2 function may induce age-related changes in acute seizure risk but disrupt integrity and connectivity of neuronal networks underlying chronic seizures. PSEN2 KO mice thus provide an untapped opportunity to assess seizures in the context of AD and aging.

The long-term consequences of acute seizures in the aged, AD-associated brain are altogether understudied. Both epilepsy and AD are defined by glutamate excitotoxicity^22; 23^. Furthermore, in some AD models, seizures worsen the related hippocampal neuropathology^24^. Thus, AD and epilepsy share hippocampal pathological similarities such that studying seizures in AD may beneficially and bi-directionally improve outcomes. Intriguingly, PSENs and APP may regulate hippocampal kainate-type glutamate receptor (KARs) subunit expression^25^. Kainic acid (KA), a KAR agonist, administration depolarizes hippocampal pyramidal cells via activation of KARs resulting in acute seizures and status epilepticus (SE)^26; 27^. Five distinct KAR subunits, GluK1-5, are expressed in the mouse hippocampus^28^. When KA is systemically delivered, it enters the brain via passive diffusion through the blood-brain barrier, achieving low concentrations^29^.

GluK5 has a high affinity for KA and thus is preferentially activated at these lower concentrations^30^. Additionally, GluK5 requires dimerization with GluK2 to incorporate into the cell membrane, thus KARs containing dimerized GluK2/GluK5 subunits likely have the largest impact on the epileptogenic action of systemically administered KA. Further, GluK2 KO mice are extremely resistant to KA-induced seizures and GluK2 KO cells exhibit a secondary loss of GluK5 ^31; 32^. KA-induced SE in rats is associated with significant age-related effects; rats ∼2 years-old or older are more sensitive to KA than younger adult rats^33; 34^. Sex differences also occur in rodent KA-SE. Aged female C57BL/6 mice are more vulnerable to KA-induced seizures than both age-matched male and younger female mice and exhibit worsened hippocampal neuronal damage and astrocytic reactivity ^35^. We thus hypothesized that hippocampal GluK2 and/or GluK5 KAR expression may differ in young and aged PSEN2 KO mice versus age-matched wild-type (WT) controls, which may underlie age-related changes in seizure susceptibility with loss of normal PSEN2 function. Our present study thus aimed to initially establish the degree to which acute seizure susceptibility changes with age in mice with and without loss of normal PSEN2 function, a known risk factor for EOAD.

## 2. Methods

### 2.1 Animals

Male and female PSEN2 KO mice were bred at the University of Washington from stock originally acquired from the Jackson Laboratory (stock #005617) as described^19; 20^. Wild-type (WT) C57BL/6J male and female control mice were purchased from Jackson Laboratory (Stock #000664) at 6-8 weeks-old and housed alongside PSEN2 KO mice until behavioral seizure testing at minimum 4-8 weeks later. Mice were housed in a 14:10 light cycle (lights on at 06h00: off at 20h00) in ventilated cages with corncob bedding and food and water provided *ad libitum*, as previously published^36^, in a manner consistent with the *Guide for the Care and Use of Laboratory Animals*. All animal work was approved by the UW Institutional Animal Care and Use Committee (protocol 4387-01) and conformed to ARRIVE guidelines^37^. All behavioral seizure testing was performed during the hours of 09h00 and 17h00.

### 2.2 Kainic Acid-induced Acute Seizures and Status Epilepticus

Young (3-4 month old; male WT = 12; female WT = 10; male PSEN2 KO = 16; female PSEN2 KO = 15) mice and aged (12-15 month old; male WT = 10; male PSEN2 KO = 12; female WT = 10; female PSEN2 KO = 11) received KA (Tocris, Cat #0222; discontinued) prepared in saline solution at a concentration of 2 mg/mL (**Figure 1**). To reduce KA-induced mortality, we opted for systemic administration of KA through repeated low doses delivered via the intraperitoneal (i.p.) route ^38; 39^. Mice received an initial 10 mg/kg KA dose, with subsequent 5 mg/kg doses every 20 minutes until SE onset (two generalized Racine stage 4+ seizures within 30 minutes), per our published protocol^40^. The dose was reduced to 2.5 mg/kg if a mouse only had a single Racine stage 4+ seizure within the 20-minute dosing window. Following SE onset, additional seizure activity was monitored and recorded for 1-hour, after which mice were given 1 mL of lactated Ringer’s solution (subcutaneous; Hospira) and allowed to recover in their home cage for the remainder of the study. Post-SE survival and body weight change from baseline (pre-SE) was recorded 1-day, 3-days, and 7-days post KA-SE. Sham-SE mice (sterile PBS every 20-minutes for 1-hour) were included as control as follows: 3-4 month old (male WT = 10; female WT = 10; male PSEN2 KO = 13; female PSEN2 KO = 12) mice and 12-15 month old (male WT = 9; male PSEN2 KO = 13; female WT = 9; female PSEN2 KO = 12). A total of 68 young mice and 51 aged mice were sacrificed 7-days post KA-SE, with brains thus collected for histology.

**Figure 1.**
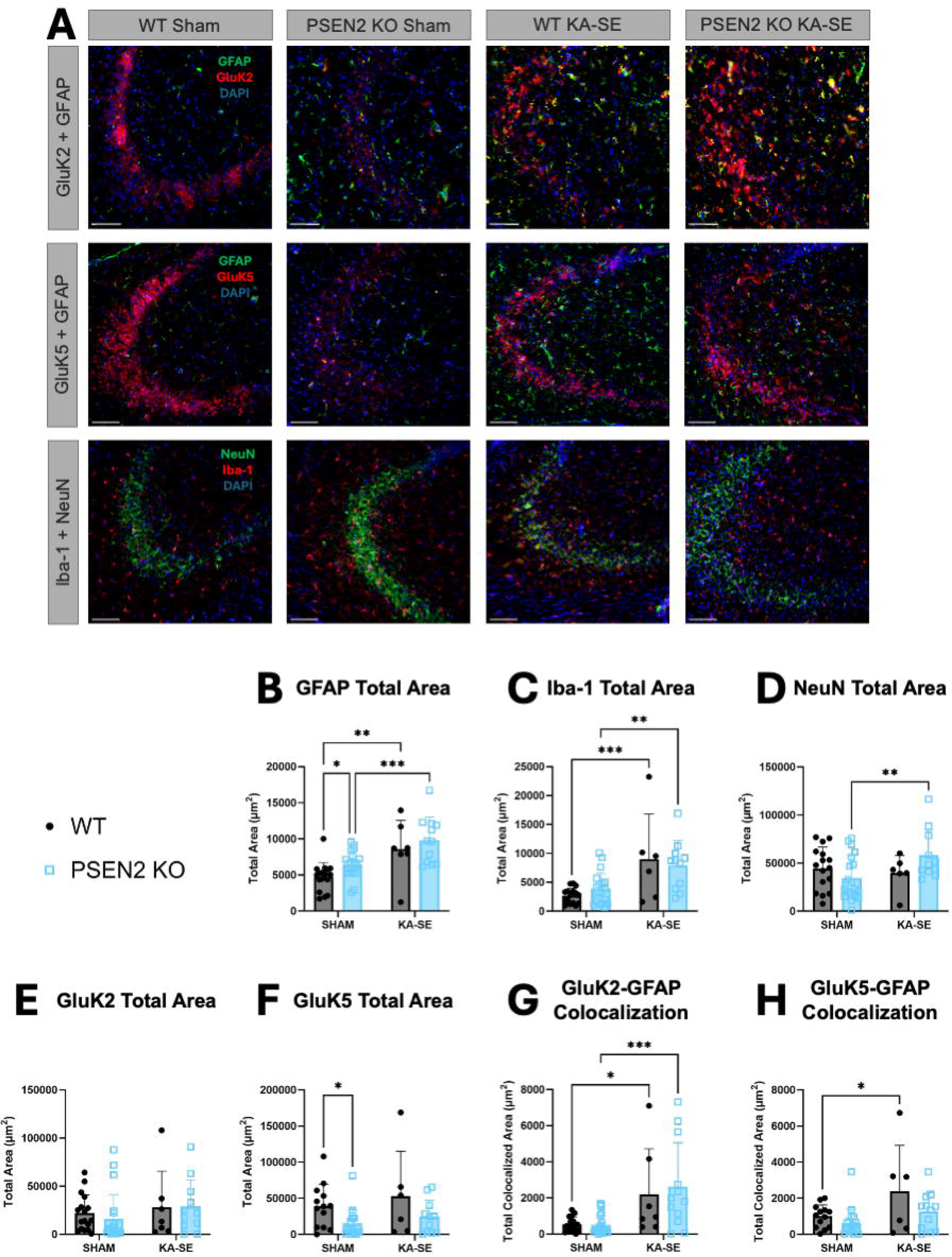
Loss of normal PSEN2 function leads to significantly reduced latency to KA-induced acute seizures and status epilepticus in both male and female young mice relative to age-matched C57BL/6J mice. A) KA-SE protocol. See Methods for details. B) 3-4-month-old male mice latency to first Racine stage 4/5 seizure (Log-rank (Mantel Cox) test X^2^ = 6.41, *p=0.011, and unpaired t-test **p = 0.003) C) 3-4-month-old male mice latency to KA-induced SE onset (Log-rank (Mantel Cox) test X^2^ = 3.14, p = 0.08, unpaired t-test *p = 0.027). D) 3-4-month old male mice amount of KA received (*p = 0.02) E) 3-4-month-old female mice latency to first stage 4/5 seizure (Log-rank (Mantel Cox) test X^2^ = 13.01, ***p = 0.0003, unpaired t-test ***p = 0.0001). F) 3-4-month-old female mice latency to SE onset (Log-rank (Mantel Cox) test X^2^ = 8.47, **p = 0.0036, unpaired t-test **p = 0.0024). G) 3-4month-old female mice amount of KA received (p value = 0.003).

### 2.3 Euthanasia

7-days after KA-SE or sham SE, all surviving mice were euthanized by live decapitation and brains collected into 4% PFA (CAT: PF101, FD NeuroTechnologies) for 24 hours. Brains were then transferred to 30% sucrose in PBS for 24-72 hours before being flash-frozen in 2-methylbutane on dry ice and stored at -80°C until sectioning on a Leica CM1850 or CM1860 cryostat into 30-µm thick coronal slices from Bregma -1.70 to -210 to capture the dorsal hippocampus for subsequent immunofluorescence. Two rostral and two caudal sections/mouse were mounted on a Superfrost slide (Fisher Scientific) and stored at -80°C until immunofluorescent processing.

### 2.4 Immunofluorescence

Cryosectioned slides were washed with 1X PBS 3 x 5 minutes before antigen retrieval. Briefly, 1X Antigen Retrieval Buffer (ab93678, Abcam) was heated to 95°C and slides were immersed in this heated buffer in the incubator for 20-minutes, then removed from the incubator and allowed to come to room temperature in the buffer for 40 minutes (60 minutes in buffer in total). Slides were then washed in dH_2_O 3 x 2 minutes before being incubated in 10% Goat Serum in 5% Triton-X PBS solution under coverwells in a humid chamber for 2 hours. Primary antibodies (NeuN (1:500; MAB377, MilliporeSigma), GFAP (1:500; MAB3402X, MilliporeSigma), GluK2 (1:300; ab66440, Abcam), GluK5 (1:500; ab67408, Abcam), Iba-1 (1:500; 019-19741, Wako)) were applied under 200 μL coverwells in a 5% Goat Serum in 1X PBS solution overnight at 4°C, with an extra 2 hour room temperature incubation for GluK5 and GluK2. The following day, coverwells were removed and slides washed in 1X PBS 3 x 10 minutes. Secondary antibodies (1:500; Goat anti-Mouse IgG Alexa Fluor 488, ab150113, Abcam; 1:500; Goat anti-Rabbit IgG Alexa Fluor 555, ab150078, Abcam) were applied under 200 μL coverwells in 1X PBS for 2 hours at room temperature then washed in 1X PBS 3 x 10 minutes. Alexa Fluor® 488 anti-β-amyloid (6E10; Biolegend Cat#803013; **Supplemental Figure 1**) histology was conducted as previously reported^24; 41^. Slides were then coverslipped with Prolong Gold with DAPI (P36935, ThermoFisher). Pictomicrographs were captured with a fluorescent microscope (Leica DM-6) with a 20x objective (80x final magnification). Acquisition settings were held constant throughout. AIVIA (Leica) cell classifier tool was used in combination with AVIA cell count tool to detect GFAP and Iba-1 expression. Signal thresholding was set using Fiji/ImageJ. A colocalization plugin was used to determine total colocalized area between GluK and GFAP expression (Credit: Pierre Bourdoncle, Institut Jacques Monod, Service Imagerie, Paris, France).

### 2.5 Statistical Analysis

Behavioral seizures were assessed using a Log-rank (Mantel Cox) test for latency to first stage 4/5 seizure and SE onset, and using t-tests for behavioral seizure duration and amount of KA administered in young and aged mice. Proportion of acute survival post-SE insult was assessed using a Fisher’s exact test. Body weight change after KA administration and SE onset was assessed in KA-treated mice using a two-way ANOVA (time x genotype), with Tukey’s post-hoc tests. Immunofluorescence data were assessed using two-way ANOVA followed by the Fisher’s least square difference post-hoc test. Parametric data (body weight, amount of KA, immunohistochemical analyses, etc) were assessed for normality using the Shapiro-Wilk test; non-parametric data (proportion tests) were analyzed accordingly. All statistical analysis was conducted using GraphPad Prism, v10.0 or later (GraphPad Software, San Diego, CA, USA), with significance defined as p < 0.05. The sample size for histological analysis in the different experimental groups was always ≥6 (mixed sex), but for behavioral studies the minimum group size was always ≥10/sex because of known differences in seizure latency between males and females.

## 3. Results

### Young PSEN2 KO mice are more susceptible to KA-induced acute seizures than age-matched wild-type mice

We first assessed the susceptibility of both WT and PSEN2 KO mice aged 3-4 months old to both KA-induced acute seizures and SE (**Figure 1A**). Young adult PSEN2 KO mice were significantly more susceptible to KA-induced seizures. Male WT mice took an average of 87.6 ± 15.5 minutes to have their first acute stage 4/5 seizure, while male PSEN2 KO mice took 61.8 ± 23.5 minutes (**Figure 1B**; X^2^=6.41, p=0.011). While the latency to KA-induced SE did not achieve statistical difference as assessed by a Log-rank test (X^2^ = 3.14, p = 0.08), the overall average time for male PSEN2 KO mice to enter SE was significantly sooner than male WT mice: 78.6 ± 25.3 minutes vs. 98.2 ± 16.0 minutes, respectively (t=2.34, p = 0.027; **Figure 1C**). Female WT mice took an average of 102 ± 16.9 minutes to have their first acute stage 4/5 seizure; female PSEN2 KO mice took significantly less time (61.3 ± 24.5 minutes - **Figure 1E**; X^2^=13.0, p=0.0003). Similarly, female PSEN2 KO mice had faster latency to SE onset (X^2^=13.0, p=0.0003) after the first KA dose, which was on average significantly sooner than their WT counterparts (80.2 ±26.4 minutes in PSEN2 KO versus 112 ± 15.1 minutes in WT mice; t=3.43, p = 0.0024 - **Figure 1F**). Once acute seizure onset was confirmed, however, there was no significant difference in average time to SE in male or female PSEN2 KO mice (**Supplemental Figure 2**). Further highlighting their increased sensitivity to KA, both male and female PSEN2 KO mice received significantly less KA than their matched WT counterparts (**Figure 1D and G**). Additionally, male PSEN2 KO had significantly greater mortality than male WT mice (**Figure 2A**), whereas genotype did not influence female mouse mortality (**Figure 2C**). Notably, 3-4-month-old WT females were significantly more susceptible to KA-induced mortality than matched WT males (**Supplemental Figure 3A**). An acute reduction in body weight immediately following KA-SE, as presently observed, is typical of this model; genotype did not influence post-SE body weight change during the 7-day monitoring period (**Figure 2B and D**).

**Figure 2.**
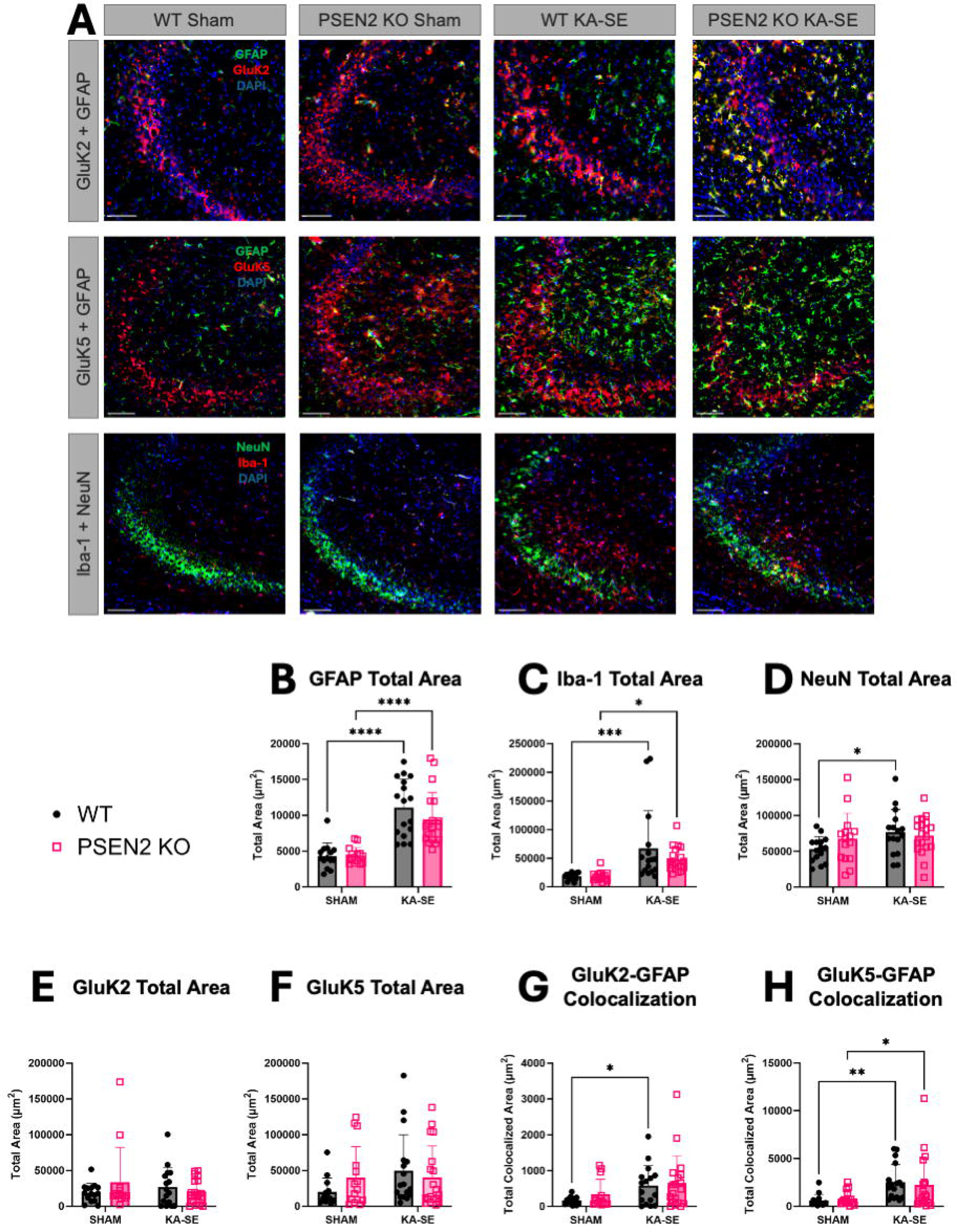
3-4-month-old male mice mortality Fisher’s exact test (p value = 0.02). B) 3-4-month-old male body weight change following KA-SE (day 0 vs day 1 WT *p value = 0.01, PSEN2 KO #p value = 0.03). C) 3-4-month-old female mortality Fisher’s exact test p value = 0.67). D) 3-4-month-old female body weight change following KA-SE (Tukey’s multiple comparison test – day 0 vs day 1 WT p value ns, PSEN2 KO p value = 0.01)

### Loss of PSEN2 function does not alter aged mouse susceptibility to KA-induced acute seizures and SE

A cohort of mice aged 12-15 months were next tested to further explore age-related differences in acute seizure susceptibility with PSEN2 loss and to address how advanced age impacts seizure vulnerability. We found that aged PSEN2 KO and WT mice had a similar latency to both first stage 4/5 seizure and SE in both sexes (**Figure 3A-D**). Additionally, when comparing KA-induced acute seizure latency between ages, we found that aged female mice of both genotypes experienced the first stage 4/5 seizure faster than their younger counterparts (**Figure 4**). Younger female WT mice took an average of 102 ± 16.9 minutes to have their first stage 4/5 seizure, while their aged counterparts took 44.7 ± 14.8 minutes (**Figure 4C**; t=8.16, p<0.0001). Similarly, aged female WT mice had faster latency to SE onset versus young WT females (112 ± 15.1 vs. 54.7 ± 14.1**; t=8.80, p<0.0001** - **Figure 4G**). Aged PSEN2 KO female mice also took less time to have their first stage 4/5 seizure (61.3 ± 24.5 vs. 43.0 ± 15.9; t=2.15, p=0.043 **- Figure 4D**) and less time to enter SE compared to young PSEN2 KO females (80.2 ±26.4 vs 53. 5 ± 13.7; t=3.05, p=0.006 **- Figure 4H**). However, no differences were observed between age groups among male WT or PSEN2 KO mice (**Figure 4**). Thus, advanced age appears to significantly increase seizure susceptibility only in female mice regardless of genotype, but the latency between first seizure and SE onset was only significantly increased in male PSEN2 KO mice (**Supplemental Figure 4**). Mortality in aged female mice of both genotypes was also remarkably high (**Figure 3H**). While there were no genotype-related differences in mortality for either sex, both aged WT and PSEN2 KO females had increased mortality compared to their younger counterparts (**Figure 4K and L**). Additionally, male WT, but not PSEN2 KO mice experienced increased age-related mortality (**Figure 4I and J**).

**Figure 3.**
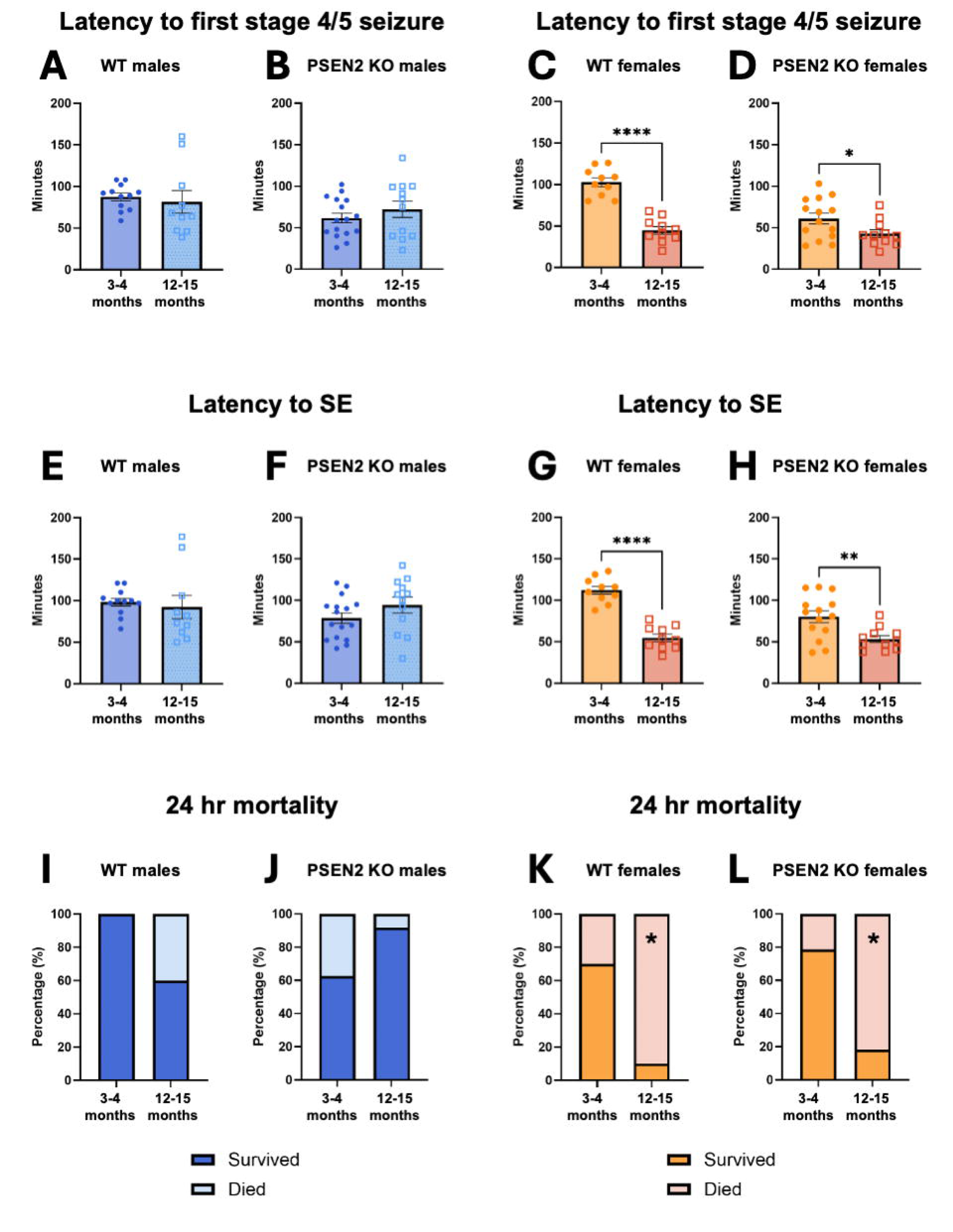
A) 12-15-month-old male mice latency to first stage 4/5 seizure (Log-rank (Mantel Cox) test X^2^ = 0.66, p value > 0.4, bar graph unpaired t-test p value > 0.5). WT male mice mean latency to first stage 4/5 seizure onset was 81.6 ± 42.8 minutes versus that of PSEN2 KO, which had a mean latency of 72.3 ± 34.5 minutes. B) 12-15-month-old female mice latency to first stage 4/5 seziure (Log-rank (Mantel Cox) test X^2^ = 0.017, p value > 0.9, and bar graph unpaired t-test p value > 0.8). The latency to first stage 4/5 seizure in WT female mice was 44.7 ± 14.8 minutes versus the latency of PSEN2 KO mean 43.0 ± 15.9 minutes. C) 12-15-month-old male mice latency to SE onset (Log-rank (Mantel Cox) test X^2^ < 0.01, p value > 0.9, and bar graph unpaired t-test p value = 0.9). The WT mean latency to SE onset was 92.40 ± 44.34 minutes versus the latency of male PSEN2 KO of 94.4 ± 33.2 minutes. D) 12-15-month-old female (Log-rank (Mantel Cox) test X^2^ < 0.01, p value > 0.9, and bar graph unpaired t-test p value > 0.8). The aged WT female mean latency to SE onset was 54.7 ± 14.0 minutes versus the PSEN2 KO mean of 53.5 ± 13.7 minutes. E) 12-15-month-old male amount of KA received (unpaired t-test p value > 0.7). F) 12-15-month-old female mice amount of KA received (unpaired t-test p value > 0.1). G) 12-15-month-old male mortality (Log-rank (Mantel-Cox) test X^2^ = 3.15, p = 0.065 and Fisher’s exact test p value = 0.1353). H) 12-15-month-old female mortality (Log-rank (Mantel-Cox) test X^2^ = 1.38, p value > 0.2401 and Fisher’s exact test p value>0.99).

**Figure 4.**
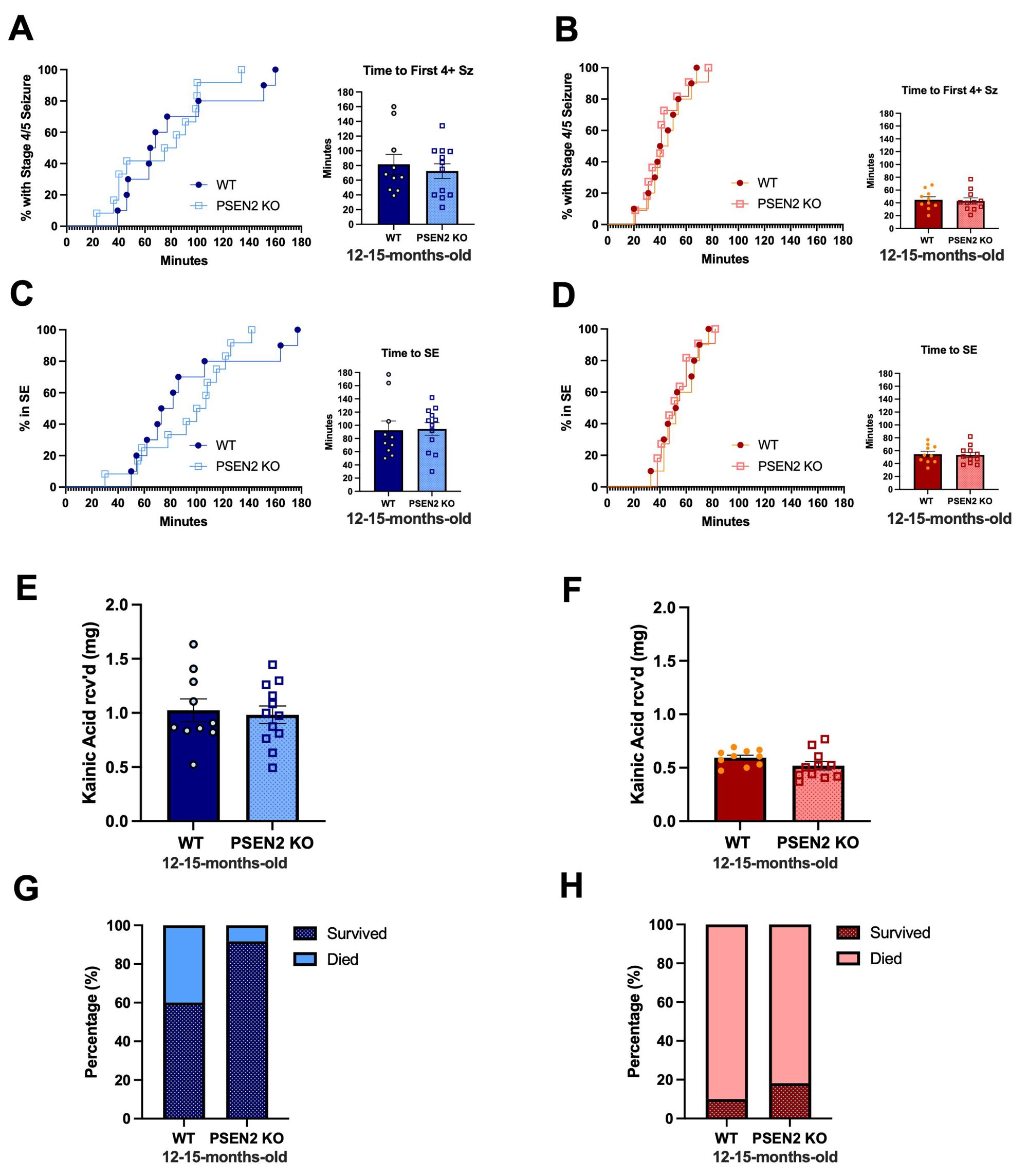
Only female mice showed worsened susceptibility to kainic acid with age. A) WT male latency to first stg 4/5 seizure (unpaired t test p>0.6). B) PSEN2 KO male latency to first stg 4/5 seizure (unpaired t test p>0.3)). C) WT female latency to first Racine stage 4/5 seizure (unpaired t test p<0.0001). D) PSEN2 KO female latency to first Racine stage 4/5 seizure (unpaired t test, p = 0.043). E) WT male latency to SE onset (unpaired t test p>0.6). F) PSEN2 KO male latency to SE onset (unpaired t test p>0.16). G) WT female latency to SE onset (unpaired t test, p<0.0001). H) PSEN2 KO female latency to SE onset (unpaired t test, p = 0.0057). I) WT male mortality (Log-rank (Mantel-Cox) test X^2^ = 5.6, p = 0.02 and Fisher’s exact test p = 0.029). J) PSEN2 KO male mortality (Log-rank (Mantel-Cox) test X^2^ = 3.1, p value = 0.08 and Fisher’s exact test p = 0.18). K) WT female mortality (Log-rank (Mantel-Cox) test X^2^ = 7.1, p = 0.008 and Fisher’s exact test (p= 0.02). L) PSEN2 KO female mortality (Log-rank (Mantel-Cox) test X^2^ = 8.6, p = 0.003 and Fisher’s exact test p = 0.005).

### Loss of PSEN2 blunts KA-SE-induced increases in hippocampal NeuN expression

As KA-SE can induce reactive astrogliosis, microgliosis, and neuronal cell death^42^, and human variants in PSEN2 have been shown to alter microglial reactivity^43^, we also examined potential differences in glial reactivity and NeuN expression, a marker of mature neuronal cell bodies^44^. Both WT and PSEN2 KO mice showed an increase in GFAP expression, a marker of reactive astrogliosis following KA-SE (F (1, 61) = 60.2, p<0.0001, **Figure 5B**). Additionally, Iba-1 expression, a marker of microglial reactivity, was increased for both genotypes following KA-SE (F (1, 61) = 20.5, p<0.0001, **Figure 5C**). This effect in glial cells was consistent in both CA1 and DG (**Supplemental Figures 7B and C; 8B and C**). Interestingly, post-hoc analysis revealed that KA-SE history only increased NeuN immunoreactivity exclusively in WT mice in hippocampal CA3 (main effect of KA history - F(1,61)=3.94, p=0.05; post-hoc p=0.024, **Figure 5D**). This KA-SE induced increase in NeuN expression was also only present in hippocampal CA1 (p = 0.006) and DG (p = 0.0005; **Supplemental Figure 7D and 8D**). Interestingly, KA-SE only induced NeuN expression in young PSEN2 KO mice in the CA3 region (p = 0.009, **Figure 6D**), an effect absent in aged WT mice. Thus, loss of normal PSEN2 function disrupts the KA-induced increase in NeuN expression 7-days after an SE insult.

**Figure 5.**
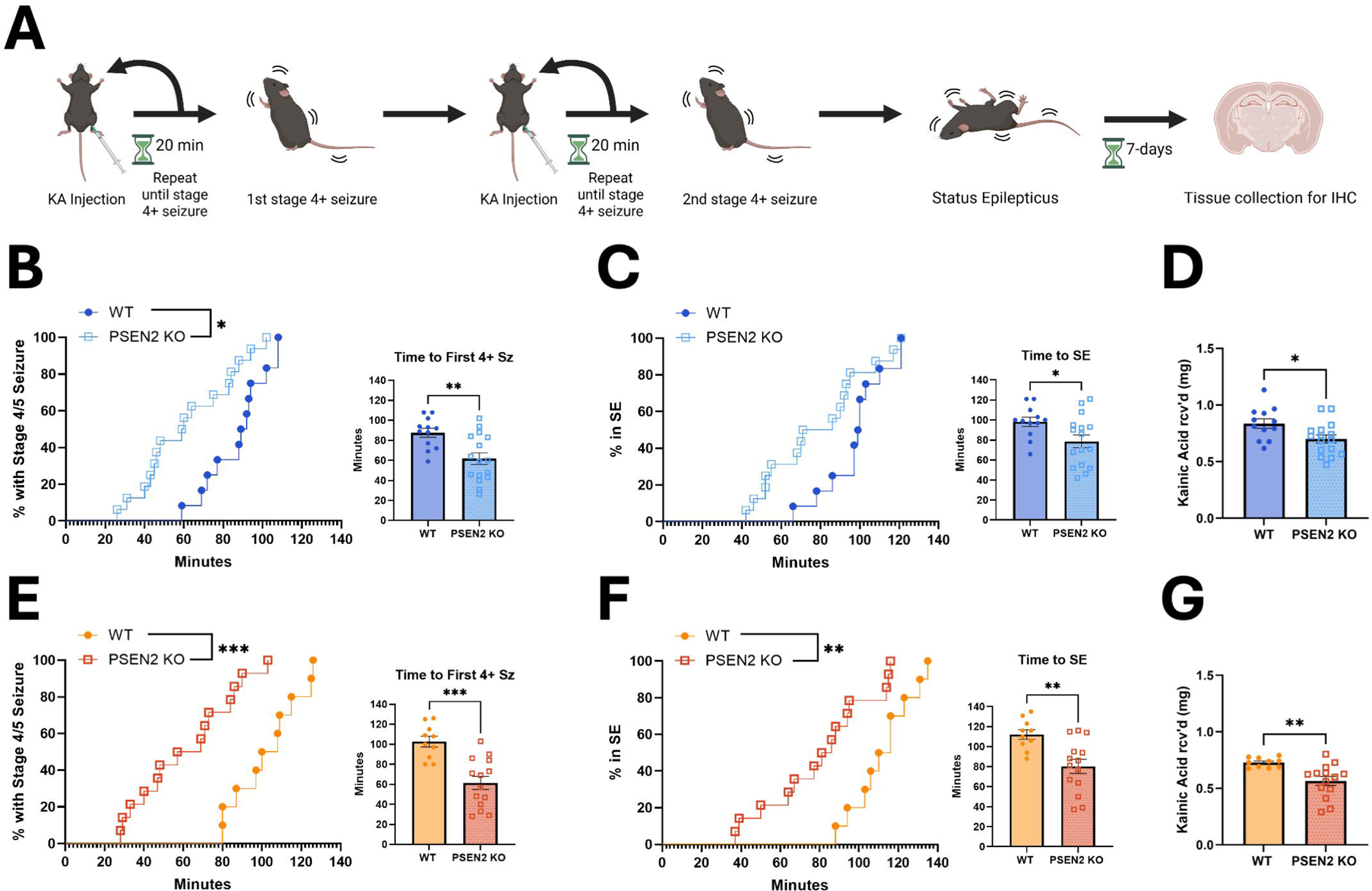
Loss of normal PSEN2 function did not impact total hippocampal expression of GluK2 and GluK5 subunits when compared to age-matched C57BL/6J mice aged 3-4 months. **A)** Representative images of GluK2 + GFAP, GluK5 + GFAP, and Iba-1 + NeuN staining at CA3 of WT and PSEN2 KO sham and KA-SE animals. **B)** GFAP expression in the CA3 region was increased in animals that received KA (Sham vs. KA-SE: F (1, 61) = 60.22, p<0.0001, WT Sham vs. KA-SE: ****p<0.0001, PSEN2 KO Sham vs. KA-SE ****p<0.0001). **C)** Iba-1 expression in the CA3 region was increased in animals that received KA (Sham vs KA-SE: F (1, 61) = 20.53, p<0.0001, WT Sham vs. KA-SE: ***p<0.0002, PSEN2 KO Sham vs. KA-SE *p<0.018). **D)** NeuN expression in the CA3 region was increased in WT animals that received KA (F (1, 61) = 3.94, p=0.05; WT Sham vs. KA-SE *p = 0.024). **E)** There was no difference in GluK2 expression in the CA3 region between either treatment groups or genotypes. **F)** There was no difference in GluK5 expression in the CA3 region between either treatment groups or genotypes. **G)** GFAP-GluK2 colocalization in the CA3 region was increased in animals that received KA (Sham vs. KA-SE: F (1, 59) = 6.91, p = 0.01, WT Sham vs. KA-SE: *p = 0.035). H) GFAP-GluK5 colocalization in the CA3 region increased in animals that received KA (F (1, 61) = 12.58, p=0.0008; WT Sham vs KA-SE **p = 0.006; PSEN2 KO Sham vs KA-SE *p = 0.036)

**Figure 6.**
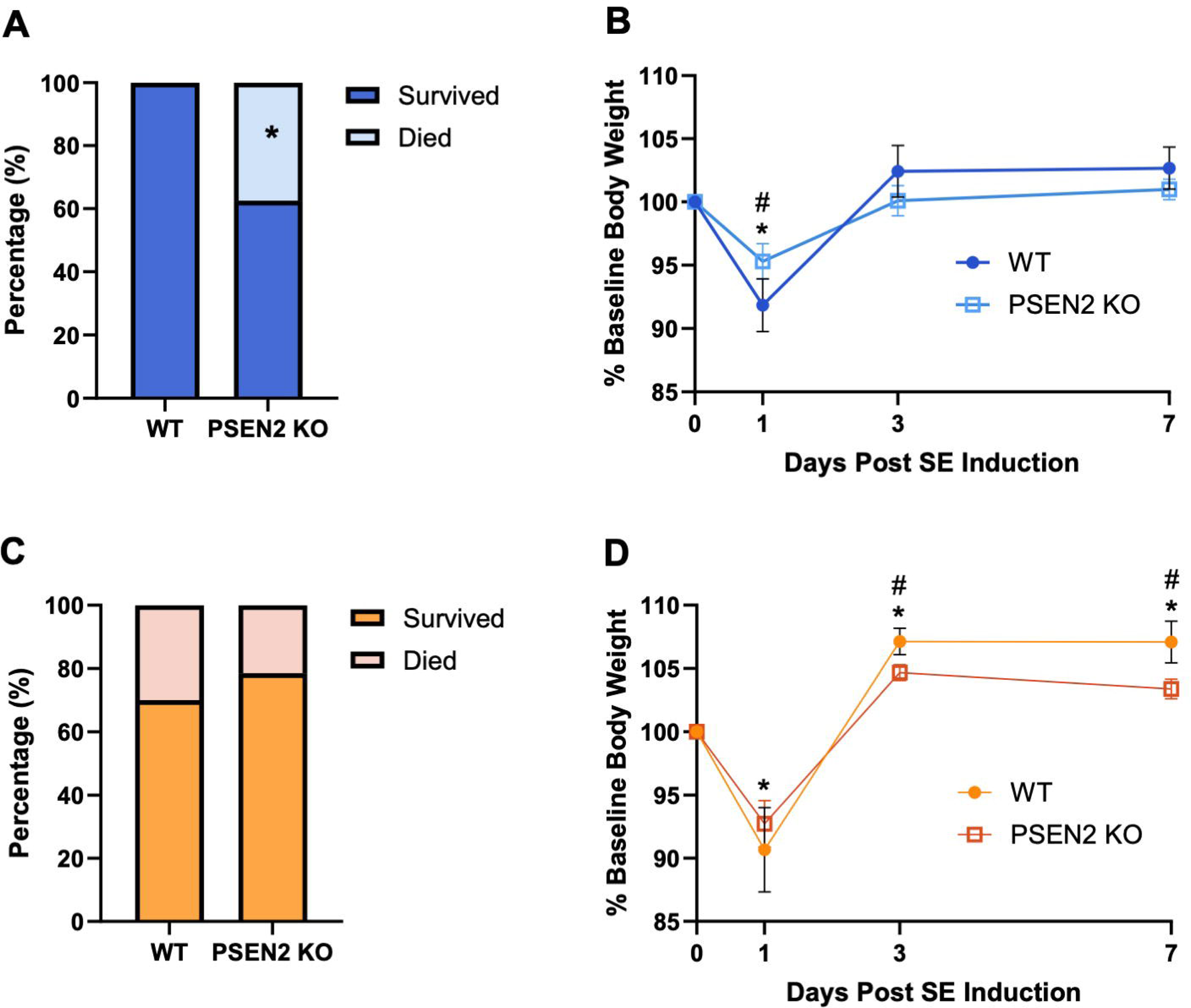
Loss of normal PSEN2 function resulted in decreased bassline expression of GluK5 compared to age-matched WT mice aged 12-15 months. **A)** Representative images of GluK2 + GFAP, GluK5 + GFAP, and Iba-1 + NeuN staining at CA3 of WT and PSEN2 KO sham and KA-SE animals. . **B)** GFAP expression in the CA3 region was increased in animals that received KA (Sham vs. KA-SE: F (1, 53) = 24.84, p<0.0001; WT Sham vs. KA-SE: **p<0.0001; PSEN2 KO Sham vs. KA-SE ***p<0.0005; Sham WT vs PSEN2 KO *p = 0.045). **C)** Iba-1 expression in the CA3 region was increased in animals that received KA (Sham vs KA-SE: F (1, 48) = 21.12, p<0.0001, WT Sham vs. KA-SE: ***p<0.0008, PSEN2 KO Sham vs. KA-SE **p<0.0058). **D)** NeuN expression in the CA3 region was increased in PSEN2 KO animals that received KA (Sham vs KA-SE: F (1, 48) = 4.121, p=0.048, PSEN2 KO Sham vs. KA-SE **p = 0.009). **E)** There was no difference in GluK2 expression in the CA3 region between either treatment groups or genotypes. **F)** PSEN2 KO mice had reduced GluK2 expression compared to WT mice (WT vs. PSEN2 KO: F (1, 45) = 7.749, p = 0.008; Sham WT vs. PSEN2 KO *p = 0.03). **G)** GFAP-GluK2 colocalization in the CA3 region was increased in animals that received KA (Sham vs. KA-SE: F (1, 53) = 20.19, p = 0.0001, WT Sham vs. KA-SE *p = 0.015; PSEN2 KO Sham vs. KA-SE p = ***0.0002). **H)** GFAP-GluK5 colocalization in the CA3 region increased in animals that received KA (Sham vs. KA-SE: F (1, 46) = 8.378, p=0.0058; WT Sham vs KA-SE *p = 0.018)

### Hippocampal KAR expression does not differ between young adult WT and PSEN2 KO mice

PSENs regulate KAR expression^25^ such that we hypothesized that a disruption in hippocampal KAR expression may underlie the increased susceptibility to KA we observed in 3-4 month-old PSEN2 KO mice. Brains of mice that survived to 7 days-post KA-SE and an additional cohort of sham mice that received saline instead of KA were examined by immunofluorescence. We quantified expression of two KAR subunits - GluK2 and GluK5 - due to their high hippocampal expression and known role in the initiation of KA-induced seizures in rodents^30; 45^. Surprisingly, we found no differences in total basal or post-SE expression of either subunit between WT and PSEN2 KO mice in the CA3 (**Figure 5E and F**). This was also the case in CA1 and DG regions (**Supplemental Figure 7E and F; 8E and F**).

### KARs are expressed on hippocampal astrocytes following KA-SE in young mice regardless of genotype

KA-SE in adult rats can induce KAR expression on astrocytes^46^. While the function of astrocytic KARs remains unknown, it has been hypothesized that they may act as a glutamate sensor for astrocytes and play a role in their reaction to excessive synaptic glutamate^46^. We were thus interested in determining if astrocytes also express KARs following KA-SE in mice and if loss of normal PSEN2 function had any effect on their expression. Therefore, we performed a colocalization analysis to assess astrocytic KAR expression (**Supplemental Figure 5**). GluK2-GFAP colocalization increased following KA-SE in CA3 (F( 1, 59) = 6.91, p = 0.01, **Figure 5G**), CA1 (F (1, 60) = 6.18, p = 0.016, **Supplemental Figure 7G**) and the DG (F (1, 60) = 4.05, p = 0.049, **Supplemental Figure 8G**). However multiple comparison analysis revealed that only WT animals had a significant increase in GluK2-GFAP colocalization following KA-SE in all three regions (CA3 - p = 0.035, CA1 - p = 0.014, DG - p = 0.041).

Similarly, KA-SE induced an increase in Gluk5-GFAP colocalization in CA3 (F (1,61) = 12.58, p = 0.0008, **Figure 5H**), CA1 (F (1,61) = 4.2, p = 0.045, **Supplemental Figure 7H**), and the DG (F (1, 62) = 7.465, p = 0.008, **Supplemental Figure 8H).** Multiple comparison analysis indicated that KA-SE led to an increase in GluK5-GFAP colocalization in the CA3 for both WT (p = 0.006) and PSEN2 KO mice (p = 0.036). Overall, both WT and PSEN2 KO demonstrated astrocytic KAR expression in hippocampal structures after KA-SE with only minimal differences between genotypes.

### Worsened microglial reactivity was associated with faster latency to first generalized acute seizure and SE solely in WT mice

We next examined how latency to first generalized stage 4/5 seizure, latency to SE, and seizure burden correlated with neuropathology (**Supplemental Figure 9**). In WT animals, there was a tendency for seizure history to be inversely correlated to Iba-1 immunoreactivity (**Supplemental Figure B**); mice that had their first stage 4/5 seizure sooner had higher Iba-1 expression in CA3 (r = -0.6134, p = 0.012) and CA1(r = -0.6102, p = 0.012). Similarly, animals that entered SE sooner tended to have higher Iba-1 expression in CA3 (r = -0.5261, p = 0.036), CA1 (r = -0.5412 , p = 0.030), and the DG (r = -0.5241, p = 0.037). These data pointed to increased microglial reactivity in WT mice that were more sensitive to KA-induced seizures.

While the amount of KA the animals received may also impact the extent of microglial reactivity, a correlation between amount of KA received and Iba-1 expression was only seen in the CA3 (r = -0.5322, p = 0.034). Interestingly, this trend was not observed in the PSEN2 KO animals.

### Loss of PSEN2 increased baseline GFAP expression in older animals

KA-SE increased GFAP and Iba-1 expression at 7 days post-insult for both genotypes in CA3 (GFAP: F (1, 53) = 24.8, p < 0.0001, Iba-1: F (1, 48) = 21.1, p < 0.0001), CA1 (GFAP: F (1, 53) = 20.6, p < 0.0001, Iba-1: F (1, 49) = 28.1, p < 0.0001), and the DG ( F (1, 53) = 13.6, p = 0.0005, Iba-1: F (1, 48) = 17.3, p = 0.0001). However, in the CA3 (p = 0.045) and DG (p = 0.048), PSEN2 KO mice had increased baseline GFAP expression compared to WT mice (**Figure 6B** and **Supplemental Figure 11B**).

### Loss of PSEN2 reduced hippocampal GluK5 expression in older animals

Despite no genotype-related difference in KA-induced acute seizure susceptibility in older mice, we observed reduced baseline GluK5 expression in aged PSEN2 KO mice versus age-matched WTs (p = 0.03, **Figure 6F**). Baseline GluK5 expression was similarly reduced in the CA1 region of PSEN2 KO mice compared to their WT counterparts (p = 0.006, **Supplemental Figure 10F**). No differences in total GluK2 expression were observed between genotypes (**Figure 6E**).

### KARs are expressed on hippocampal astrocytes following KA-SE in aged WT and PSEN2 KO mice

GluK2-GFAP colocalization increased following KA-SE in CA3 (F (1, 53) = 20.19, p < 0.0001, **Figure 6G**), CA1 (F (1, 53) = 20.6, p < 0.0001, **Supplemental Figure 10G**) and the DG (F (1, 53) = 13.8, p = 0.0005, **Supplemental Figure 11G**). In the CA3 and CA1 regions, both WT (CA3 p = 0.015, CA1 p = 0.005) and PSEN2 KO (CA3 p = 0.0002, CA1 p = 0.0007) mice had increased astrocytic GluK2 expression 7-days after KA-SE (**Figure 6G and Supplemental Figure 10G**). However, astrocytic GluK2 expression was significantly increased in DG of only PSEN2 KO mice (p = 0.0006, **Supplemental Figure 11G**). GluK5-GFAP colocalization increased following KA-SE in CA3 region (F (1, 46) = 8.38, p = 0.006, **Figure 6H**). Astrocytic GluK5 expression increased following KA-SE in WT mice only in the CA3 region (p = 0.0178, **Figure 6H**).

### Worsened seizure burden was only associated with increased astrogliosis in PSEN2 KO mice

In PSEN2 KO mice, there was a tendency for GFAP expression to increase with seizure burden in CA3 (r = 0.8, p = 0.04) and CA1(r = 0.7, p = 0.03, **Supplemental Figure 12B**). Among aged WT mice, higher GluK2 expression in CA1 was associated with increased latency to both first Racine stage 4/5 seizure (r = 0.9, p = 0.008) and SE (r = 0.9, p = 0.005), as well as a higher amount of administered KA (r = 0.9, p = 0.005, **Supplemental Figure 12C**). These data likely show that larger amounts of systemically-delivered KA are associated with increased hippocampal GluK2 expression 7-days later.

## 4. Discussion

Our present study significantly extends and expands our earlier work to assess the age-related seizure susceptibility associated with loss of normal PSEN2 function^19; 20^. It carries several important findings to better understand the intersection between seizures and AD in the context of PSEN2 deficiency. Most importantly, this study suggests that loss of normal PSEN2 function accelerates senescence, promoting greater seizure vulnerability earlier than unaffected individuals. This finding is particularly compelling considering that PSEN2 KO mice do not show Aβ accumulation (**Supplemental Figure 1** and ^12^), but are primed for neuroinflammation ^13; 14^, allowing for the investigation of Aβ-independent drivers of seizure risk in the AD-associated and aged brain. First, we demonstrate that young adult PSEN2 KO mice have increased KA-SE susceptibility. PSEN2 KO mice had both reduced latency to their first generalized tonic-clonic seizure and SE compared to age-matched WT mice. These data align well with our previous findings that 2–4-month-old PSEN2 KO mice have a reduced minimal clonic threshold, a model of acute forebrain focal seizures, compared to age-matched WT mice^20^. Secondly, this increased susceptibility to KA-induced acute seizures and SE disappears in aged mice, as mice >12-15-months-old had similar KA-SE susceptibility regardless of genotype. These data also align with our previous findings, as both WT and PSEN2 KO mice aged 8-months-old had a similar minimal clonic threshold. Taken together, these present data show that young adult PSEN2 KO mice are more prone to acute seizures versus age-matched WT mice. However, we have demonstrated previously that young adult PSEN2 KO mice seem to be resistant to formation of a hyperexcitable neuronal network via corneal kindling^19; 20^. This resistance to corneal kindling is also age-dependent, as older PSEN2 KO and WT mice have a similar rate of kindling acquisition, suggesting that PSEN2 function influences neuronal hyperexcitability in an age-dependent manner. Additionally, here we determined that aged female mice are more susceptible to KA than young adult female mice, regardless of genotype. Another study looking into sex differences in the KA-SE model reported no differences in seizure susceptibility or mortality between males and females (2-3 months). However, this same study also found that gonadectomized mice were more susceptible to generalized seizures and SE compared to same-sex control mice. This effect as well as seizure-induced mortality was also significantly increased in gonadectomized males^47^. Our present study also found a dramatic increase in mortality among aged females of both genotypes, pointing to a potential synergistic effect of age and sex on susceptibility to KA-SE. These differences in KA-SE susceptibility across age groups in the WT female mice may be due to changes in the expression of PSENs during normal aging. Previous studies have shown that PSEN1 expression decreases in the brains of aged mice whereas PSEN2 expression increases^48^. Further studies will need to be done to determine how changes in PSEN1 expression due to loss of normal PSEN2 function impact seizure susceptibility.

Our study demonstrates that both age and AD genotype can influence KAR subunit expression differences. While KARs typically act in an ionotropic manner following activation with an agonist such as KA or glutamate, GluK5 subunits may mediate inhibition of post-depolarization hyperpolarizing current through metabotropic mechanisms^49^. Mossy fiber KARs containing GluK2 and GluK3 act as autoreceptors that facilitate synaptic transmission and long and short-term plasticity^50; 51^. While KARs are known to play a role in epilepsy, how KARs regulate hippocampal neuronal activity with advanced age is still less clear. Importantly, PSEN and APP are known to regulate hippocampal KAR levels^25^. Thus, our study had the secondary objective of determining how advanced age and loss of normal PSEN2 function influenced KAR expression prior to and after an acutely evoked seizure insult to define whether these receptors may influence subsequent epileptogenesis. We did not observe any morphological changes in young adult mice that could help explain the differences we see in KA-SE susceptibility across genotypes. Nonetheless, GluK5 expression was markedly lower in multiple hippocampal regions exclusively in aged PSEN2 KO mice. Barthet et al previously demonstrated that loss of both PSEN1 and PSEN2 in animals aged 5 to 8 weeks results in a decrease in hippocampal GluK2 expression, which is accompanied by decreased KAR-EPSCs that may contribute to cognitive decline. Additionally, they found that GluK2 interacts with APP, suggesting that PSENs may regulate KAR expression through their cleavage of APP^25^. Taken together, these data show that PSEN2 may become more important for the regulation of KARs in aging and late life.

Astrocytes express KARs after both the seizure-free latent period and later during the onset of spontaneous seizures in the rat post-SE model^46^, but the expression of KARs in mouse AD models has thus far not been robustly assessed. Here, we report that both WT C57BL/6J and PSEN2 KO mice also exhibit KAR colocalization on astrocytes 7 days after KA-induced SE. We observed astrocytic KAR expression in both WT and PSEN2 KO mice, however, the extent of expression was age-, genotype-, and hippocampal brain region-dependent. As this is only the second study to describe astrocytic KARs and these receptors do not appear to be expressed under basal conditions, very little is known about their function in health and disease. Vargas et al. hypothesized that these astrocytic KARs could either aid astrocytes in protecting neurons in excitotoxic conditions or further contribute to hyperexcitability through release of their own gliotransmitters due to increases in intracellular Ca^2+^ via activation of astrocytic glutamate receptors^46; 52–56^. Additionally, KA-SE is known to induce reactive astrogliosis that is highly linked with epilepsy ^57^. While we did not observe a difference in astrogliosis between genotypes in the young adult animals, older PSEN2 KO mice presented with exaggerated reactive astrogliosis at baseline compared to WT mice. Previous studies have found that PSEN2 variant-derived astrocytes express more GFAP than WT controls and both PSEN2 variant-derived astrocytes and microglia show exaggerated secretion of pro-inflammatory cytokines^16^. Thus, loss of normal PSEN2 function may induce a ‘primed’, pro-inflammatory state. This primed state may act as a “first hit” that could further accelerate AD pathology when challenged by a secondary insult^58^. Future histological studies must thus further investigate how KA-induced acute seizures influence pathological hallmarks of AD, such as Aβ and p-tau, across the lifespan.

While KA-SE is known to evoke neuronal death, this is highly dependent upon the model organism^42^. C57BL/6 mice, as presently utilized, has been shown to be resistant to KA-induced neurodegeneration^59^. In line with this, we did not observe any neurodegeneration following KA-SE in this study as assessed by Fluoro-Jade C staining (data not shown). In fact, we observed a consistent increase in NeuN expression across multiple regions of the hippocampus in young adult WT mice and aged PSEN2 KO mice. Neurogenesis following KA-SE has been reported previously in rat models and contributes to formation of an epileptogenic network^60; 61^. However, seizure-induced neurogenesis has only been observed in dentate granule cells^60^. Interestingly, this increase in NeuN was not observed in young adult PSEN2 KO mice. Furthermore, among aged animals, PSEN2 KO mice showed an increase in NeuN expression following KA-SE, whereas WT mice did not. Both AD and epilepsy have been shown to impact NeuN expression independently of neuronal cell count. In both AD mouse models and human samples, NeuN immunoreactivity decreases in the hippocampus^44; 62; 63^. Additional studies are needed to determine the cause of these changes in NeuN expression following KA-SE.

Overall, we demonstrate that loss of normal PSEN2 function may drive preponderance for seizures by promoting an accelerated senescence phenotype in the adult brain. Our study is the first to demonstrate that PSEN2 KO mice are more susceptible to KA SE, despite lack of accumulated Aβ, suggesting that additional factors may increase seizure vulnerability in AD. Loss of PSEN2 increases susceptibility to KA-induced seizures early in life, matching rates of aged mice, whereas this difference is lost in late life. We also show that loss of PSEN2 later in life leads to exaggerated astrogliosis, which has previously been associated with aging, further supporting the hypothesis that neuroinflammation may increase neuronal hyperexcitability in AD, which can indirectly accelerate neuropathological damage. While the present shifts in seizure susceptibility could not be explained by hippocampal KAR expression differences, loss of GluK5 expression in older PSEN2 KO mice may reflect an increased reliance on PSEN2 to regulate KAR expression with advanced age. We also observed an increase in NeuN expression following KA-SE only in WT animals, which could be indicative of differences in post-seizure neurogenesis due to loss of PSEN2. Altogether, this study demonstrates that PSEN2 is a unique and understudied contributor to AD-related seizure risk, with seizures being known to accelerate functional and pathological decline in AD.

