## Supplemental Figures for "Loss of presenilin 2 function age-dependently increases susceptibility to kainate-induced acute seizures and blunts hippocampal kainate-type glutamate receptor expression"

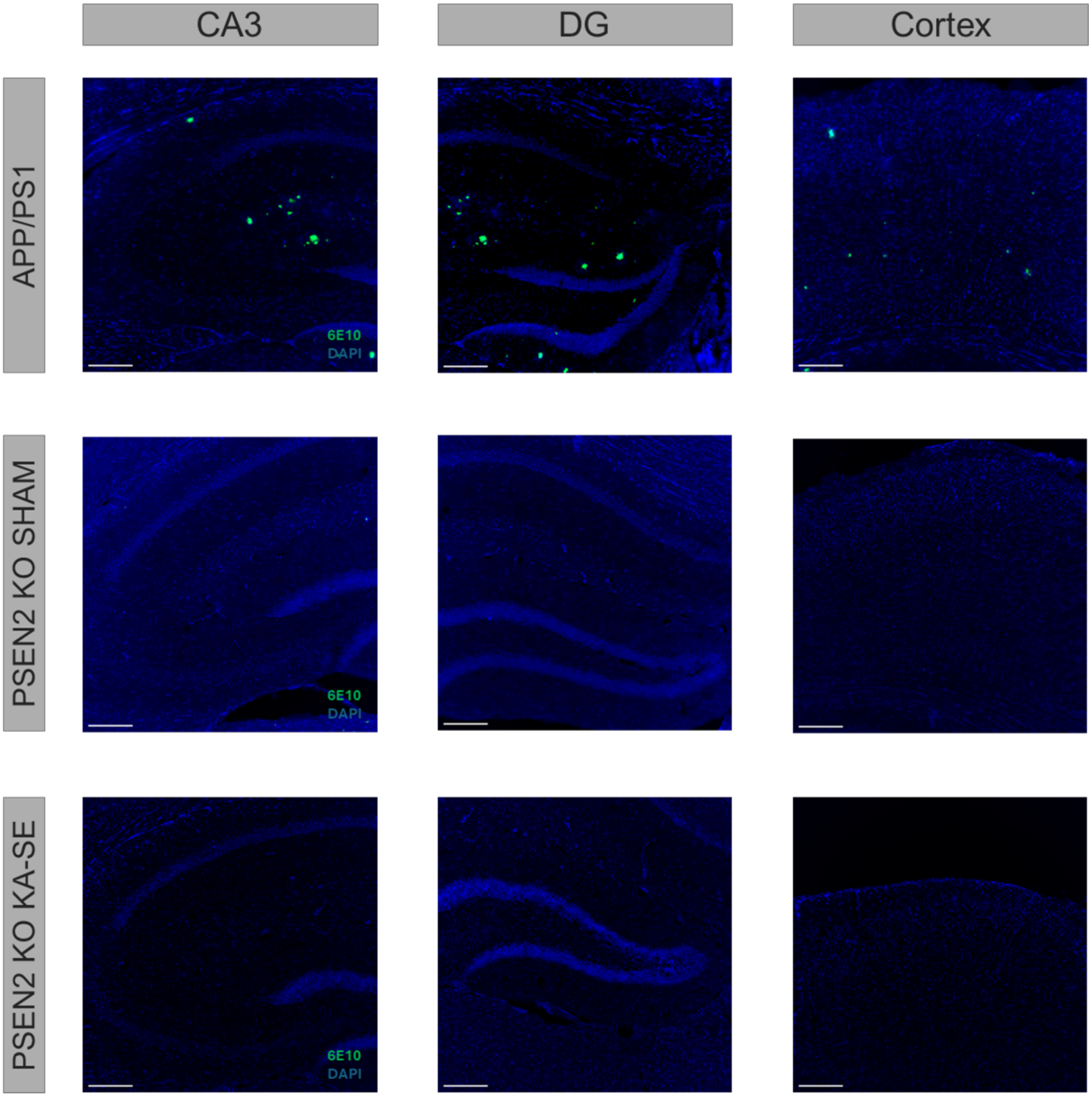


**Supplemental Figure 1. 6E10 Staining in aged PSEN2 KO mice reveals no detectable amyloid-β plaque accumulation, unlike in aged APP/PS1 mice.** IHC labeling with Alexa Fluor® 488 anti-β-amyloid (6E10; Biolegend Cat#803013) in the hippocampus and cortex of 10-month-old APP/PS1 mice and 12-15-month-old PSEN2 KO mice that received either kainic acid (KA-SE) or saline (Sham). No 6E10-positive signal was present in aged PSEN2 KO mice of either treatment group. Blue is DAPI nuclear counterstain.

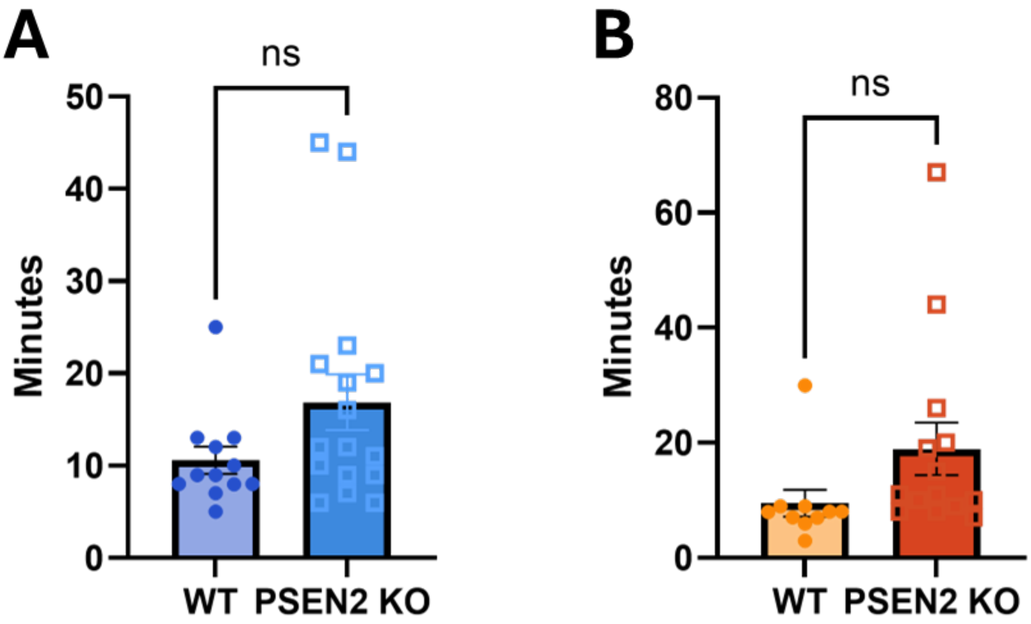


**Supplemental Figure 2 – Loss of normal PSEN2 function does not significantly change the latency between onset of first stage 4/5 seizure and SE onset in young mice.** There was no significant difference in the delay between first Racine scale stage 4/5 seizure and SE onset between genotypes in young **A)** male and **B)** female mice.

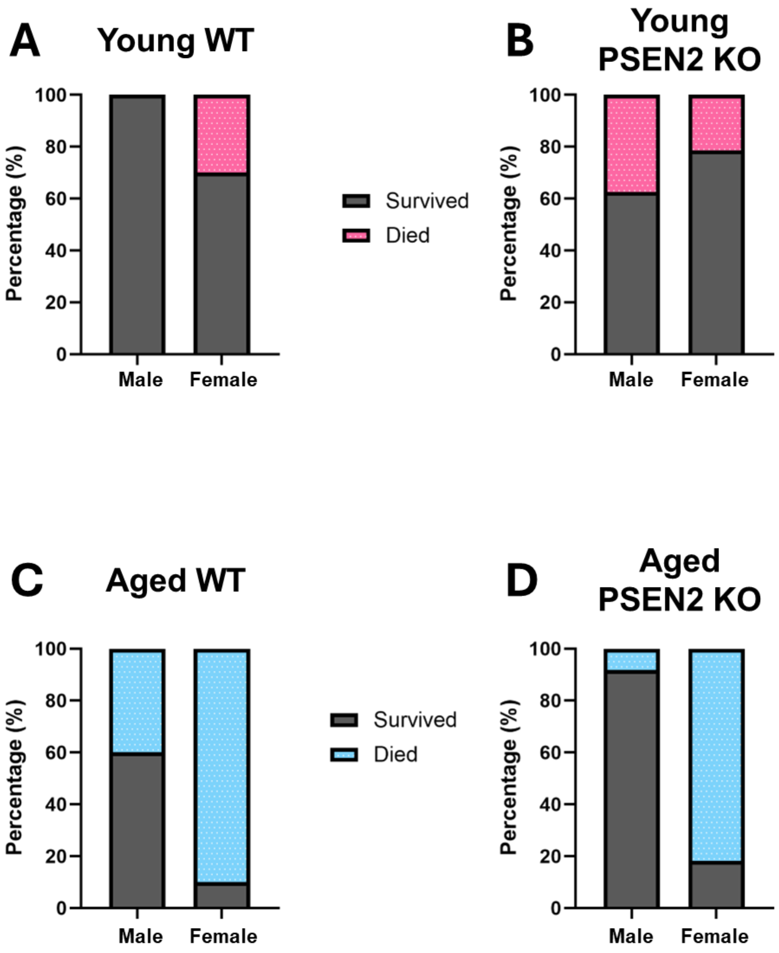


**Supplemental Figure 3 – Age and biological sex differentially influence acute survival after KA administration in mice with loss of normal PSEN2 function. A)** Young WT females did not experience significant differences in mortality from young WT males when assessed with Fisher's exact test ($p = 0.078$). **B)** There were no differences in mortality between sexes in young PSEN2 KO mice as measured by a Fisher's exact test ($p = 0.44$). **C)** In aged mice, female WT had trend for significantly greater mortality compared to males as assessed by Fisher's exact test ($p = 0.057$). **D)** Aged PSEN2 KO females had increased mortality compared to males as assessed by a Fisher's exact test ($p = 0.0006$).

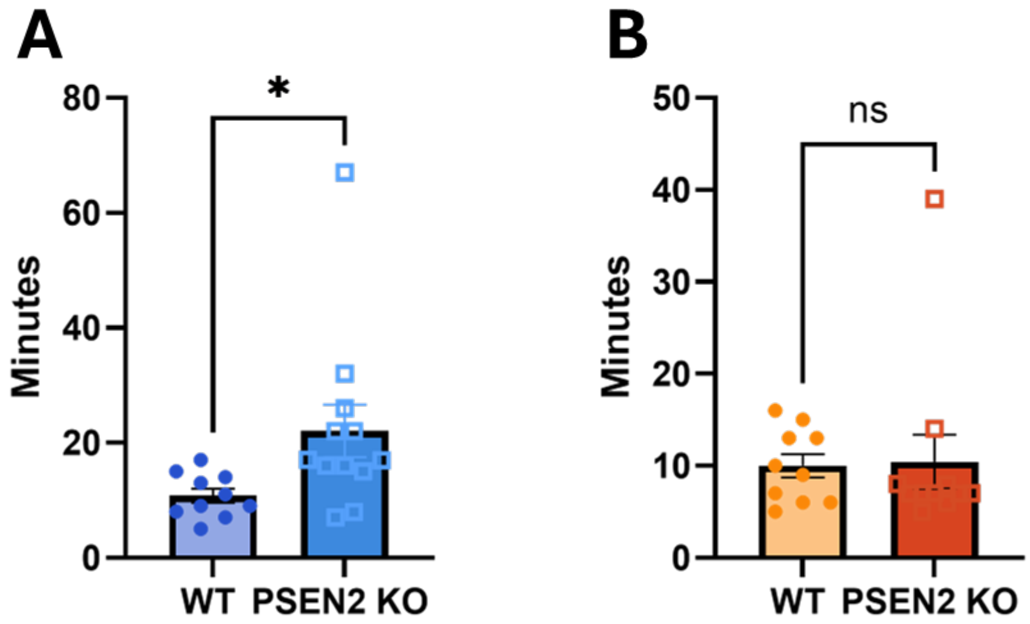


**Supplemental Figure 4 – Loss of normal PSEN2 function does significantly increase the time between first stage 4/5 Seizure and SE onset in aged male, but not female, mice.** **A)** Aged male PSEN2 KO mice took significantly longer to enter SE after their first Racine scale stage 4/5 seizure compared to age-matched WT mice. **B)** There was no significant difference in delay between first stage 4/5 seizure and SE onset between genotypes in aged female mice.

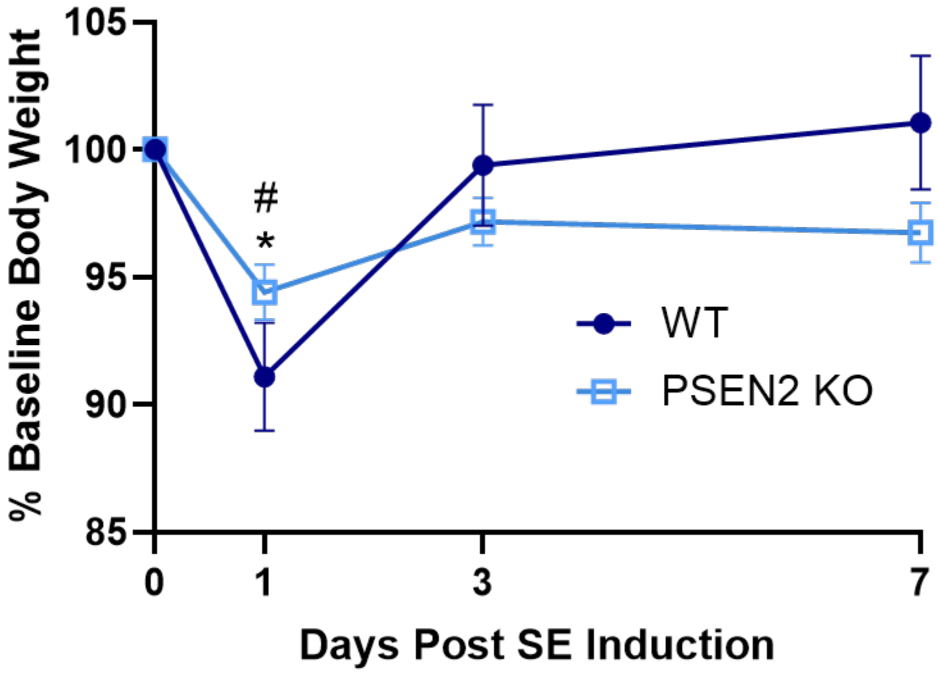


**Supplemental Figure 5 – Aged male body weight change is not significantly different between WT and PSEN2 KO mice for the 7 days following an acute KA-SE insult.** There was no effect of genotype on aged male body weight change in the week following KA-SE. Both WT and PSEN2 KO mice experienced weight loss 24 hours after KA-SE (Day 0 vs Day WT $p = 0.05$, PSEN2 KO $p = 0.0026$), but there were no significant changes in body weight over the following 7-day monitoring period as measured by a 2-way ANOVA.

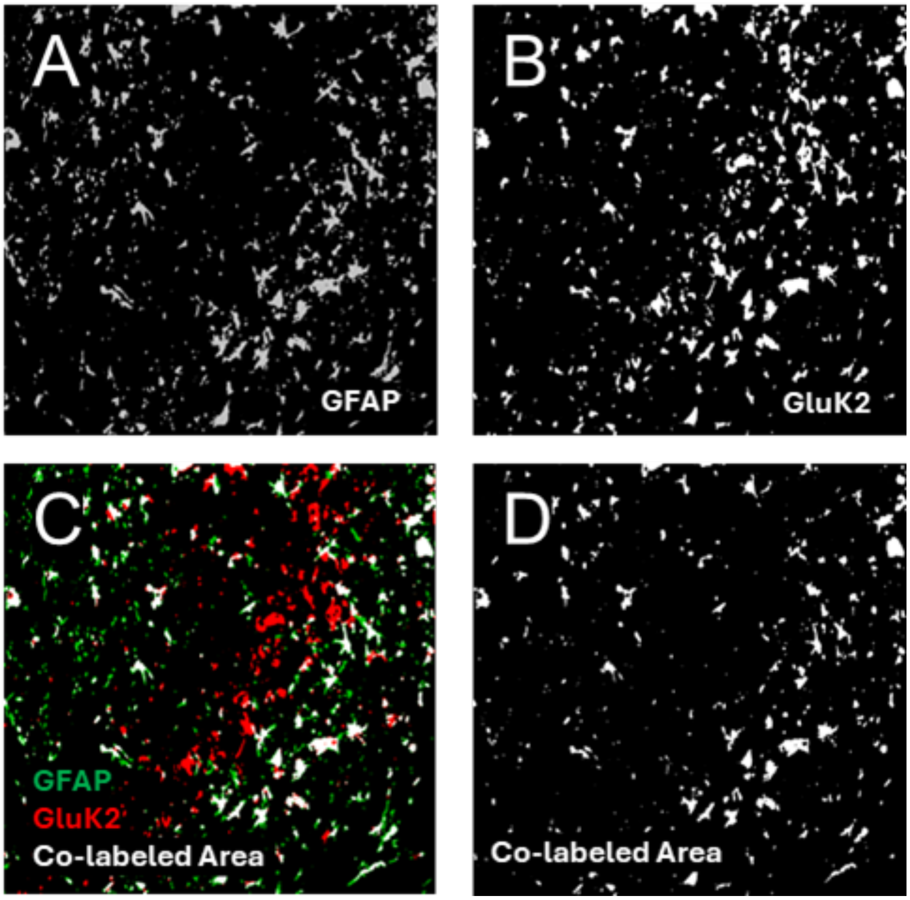


**Supplemental Figure 6 – Sample Image J colocalization workflow.** Demonstration of colocalization analysis. **A)** GFAP staining in the CA3 region 7 days post-KA-SE in example PSEN2 KO animal. **B)** GluK2 staining in the CA3 region 7-days post KA-SE in example PSEN2 KO animal. **C)** An overlay of GFAP (green) and GluK2 (red) with co-labeled areas in white. **D)** GFAP-GluK2 co-labeled area. Co-labeled images such as this one were measured for total area. This number was then used in subsequent analyses.

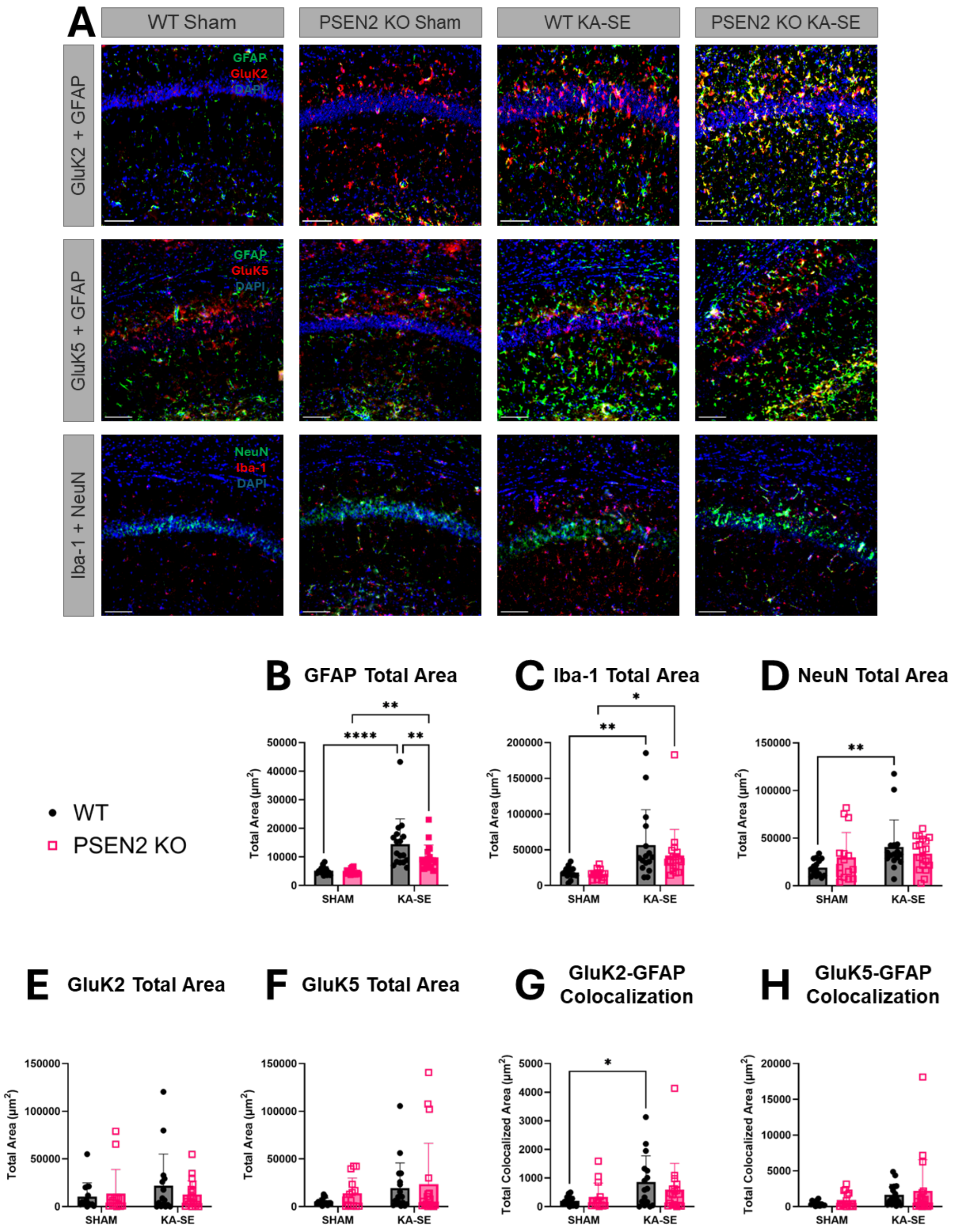
A
WT Sham
PSEN2 KO Sham
WT KA-SE
PSEN2 KO KA-SE
GluK2 + GFAP
GFAP
GluK2
DAPI
GluK5 + GFAP
GFAP
GluK5
DAPI
Iba-1 + NeuN
NeuN
Iba-1
DAPI
WT
PSEN2 KO
B GFAP Total Area
C Iba-1 Total Area
D NeuN Total Area
E GluK2 Total Area
F GluK5 Total Area
G GluK2-GFAP Colocalization
H GluK5-GFAP Colocalization
Total Area (μm²)
Total Colocalized Area (μm²)
SHAM
KA-SE

**Supplemental Figure 7 – Young mouse histopathological changes in area CA1 of dorsal hippocampus.** Loss of normal PSEN2 function did not impact total hippocampal expression of GluK2 and GluK5 subunits when compared to age-matched C57BL/6J mice aged 3-4 months. **A)** Representative images of GluK2 + GFAP, GluK5 + GFAP, and Iba-1 + NeuN staining at CA1 of WT and PSEN2 KO sham and KA-SE animals. **B)** GFAP expression in the CA1 region was increased in animals that received KA (Sham vs. KA-SE: $F (1, 62) = 31.8$, $p<0.0001$; WT Sham vs. KA-SE: ****$p<0.0001$; PSEN2 KO Sham vs. KA-SE **$p = 0.006$; KA-SE WT vs. PSEN2 KO **$p = 0.008$). **C)** Iba-1 expression in the CA1 region was increased in animals that received KA (Sham vs KA-SE: $F (1, 61) = 16.3$, $p = 0.0002$; WT Sham vs. KA-SE: **$p = 0.002$; PSEN2 KO Sham vs. KA-SE *$p = 0.021$). **D)** NeuN expression in the CA1 region was increased in WT animals that received KA (Sham vs. KA-SE: $F (1, 61) = 5.84$, $p = 0.019$; WT Sham vs. KA-SE **$p = 0.006$). **E)** There was no difference in GluK2 expression in the CA1 region between either treatment groups or genotypes. **F)** There was no difference in GluK5 expression in the CA1 region between either treatment groups or genotypes. **G)** GFAP-GluK2 colocalization in the CA3 region was increased in animals that received KA (Sham vs. KA-SE: $F (1, 60) = 6.18$, $p = 0.016$; WT Sham vs. KA-SE: *$p = 0.014$). **H)** GFAP-GluK5 colocalization in the CA3 region increased in animals that received KA (Sham vs. KA-SE: $F (1, 62) = 4.20$, $p = 0.045$)

Robinson-Cooper, Davidson et al.
*Loss of presenilin 2 function age-dependently increases susceptibility to kainate-induced acute seizures and blunts hippocampal kainate-type glutamate receptor expression*
Supplemental Figures

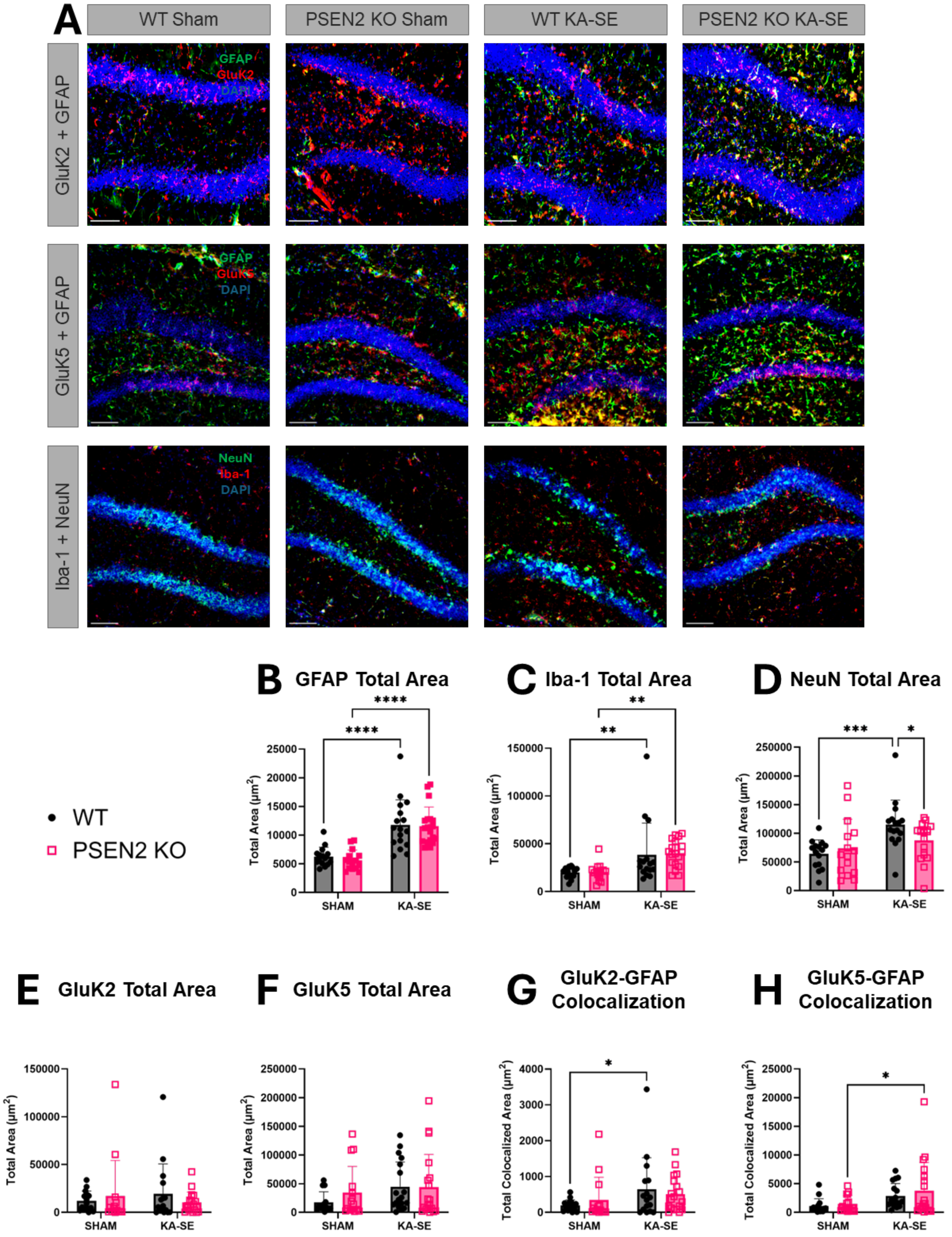

**Supplemental Figure 8 – Young mouse histopathological changes in dentate gyrus (DG) of dorsal hippocampus**. Loss of normal PSEN2 function did not impact total hippocampal expression of GluK2 and GluK5 subunits when compared to age-matched C57BL/6J mice aged 3-4 months. **A)** Representative images of GluK2 + GFAP, GluK5 + GFAP, and Iba-1 + NeuN staining at DG of WT and PSEN2 KO sham and KA-SE animals. **B)** GFAP expression in the DG was increased in animals that received KA (Sham vs. KA-SE: $F(1, 62) = 56.4$, $p<0.0001$; WT Sham vs. KA-SE: ****$p<0.0001$; PSEN2 KO Sham vs. KA-SE **** $p<0.0001$). **C)** Iba-1 expression in the DG was increased in animals that received KA (Sham vs KA-SE: $F(1, 61) = 16.6$, $p = 0.0001$; WT Sham vs. KA-SE: **$p = 0.009$; PSEN2 KO Sham vs. KA-SE **$p = 0.003$). **D)** NeuN expression in the DG was increased in WT animals that received KA (Sham vs KA-SE: $F(1, 61) = 11.3$, $p = 0.001$; WT Sham vs. KA-SE ***$p = 0.0005$; KA-SE WT vs. PSEN2 KO *$p = 0.038$). **E)** There was no difference in GluK2 expression in the DG between either treatment groups or genotypes. **F)** There was no difference in GluK5 expression in the DG between either treatment groups or genotypes. **G)** GFAP-GluK2 colocalization in the DG was increased in animals that received KA (Sham vs. KA-SE: $F(1, 62) = 4.20$, $p = 0.049$; WT Sham vs. KA-SE: *$p = 0.041$). **H)** GFAP-GluK5 colocalization in the DG increased in animals that received KA ($F(1, 62) = 7.47$, $p = 0.008$; PSEN2 KO Sham vs. KA-SE: *$p = 0.030$)

Robinson-Cooper, Davidson et al.
*Loss of presenilin 2 function age-dependently increases susceptibility to kainate-induced acute seizures and blunts hippocampal kainate-type glutamate receptor expression*
Supplemental Figures

**A** 3–4-month r-value Table

| Acute Seizure Data | | CA3 | | | | | CA1 | | | | | Dentate Gyrus | | | | |
|---|---|---|---|---|---|---|---|---|---|---|---|---|---|---|---|---|
| | | GFAP | GluK5 | GluK2 | NeuN | Iba-1 | GFAP | GluK5 | GluK2 | NeuN | Iba-1 | GFAP | GluK5 | GluK2 | NeuN | Iba-1 |
| WT | KA Received | -0.02 | -0.05 | 0.01 | 0.12 | **-0.53** | -0.08 | -0.07 | -0.2 | 0.06 | -0.5 | -0.04 | 0.06 | -0.05 | 0.02 | -0.5 |
| | Time to 1st GTC | -0.3 | 0.4 | 0.05 | 0.1 | **-0.6** | -0.2 | 0.4 | -0.2 | -0.1 | **-0.6** | -0.3 | 0.4 | -0.02 | -0.08 | -0.5 |
| | Time to Status Epilepticus | -0.3 | 0.4 | 0.1 | 0.07 | **-0.5** | -0.2 | 0.4 | -0.1 | -0.1 | **-0.5** | -0.2 | 0.3 | -0.03 | -0.02 | **-0.5** |
| | Seizure Burden | 0.03 | 0.06 | -0.2 | -0.3 | 0.3 | 0.1 | 0.2 | -0.03 | -0.004 | 0.3 | 0.3 | 0.1 | -0.2 | -0.3 | 0.1 |
| PSEN2 KO | KA Received | -0.1 | -0.3 | 0.1 | -0.3 | -0.1 | -0.2 | -0.2 | -0.05 | -0.4 | -0.2 | -0.2 | -0.2 | 0.1 | -0.4 | -0.3 |
| | Time to 1st GTC | -0.002 | -0.4 | -0.1 | -0.4 | -0.2 | -0.1 | -0.2 | -0.2 | **-0.5** | -0.2 | -0.02 | -0.2 | -0.3 | -0.4 | -0.4 |
| | Time to Status Epilepticus | -0.1 | -0.4 | 0.004 | -0.24 | -0.2 | -0.2 | -0.3 | -0.3 | -0.4 | -0.3 | -0.07 | -0.3 | -0.1 | -0.4 | -0.2 |
| | Seizure Burden | 0.2 | 0.06 | -0.008 | -0.03 | 0.3 | 0.3 | 0.1 | 0.1 | 0.09 | 0.2 | -0.02 | 0.025 | 0.2 | 0.2 | 0.1 |

**B** Iba-1 Expression vs Acute Seizure Data Correlation

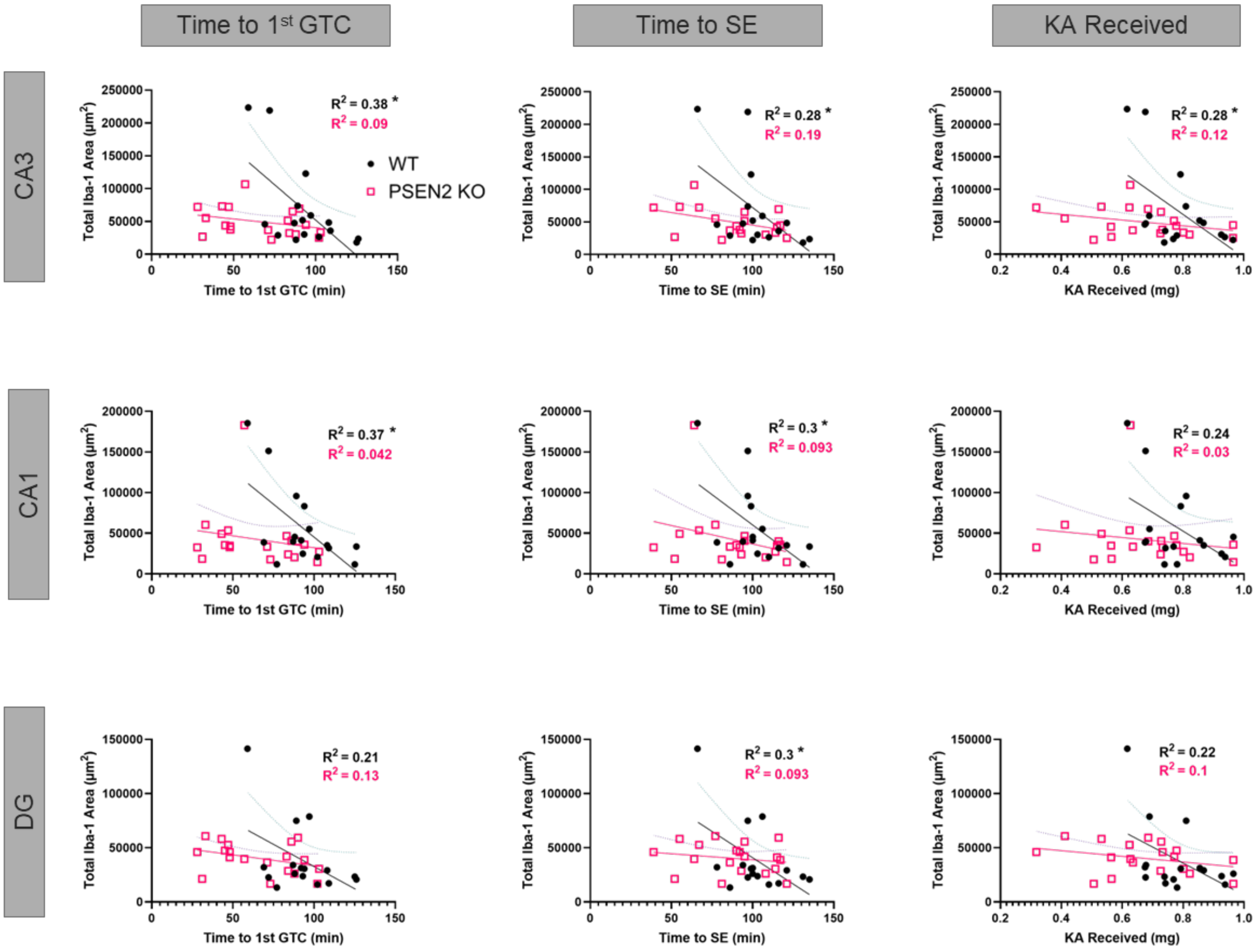


**Supplemental Figure 9 – Acute seizure burden correlations to post-KA SE histology in young mice. A)** table of r-values obtained from each analysis. Red cells denote a significant correlation. **B)** Scatterplots of Total Iba-1 expression over time to 1st GTC, time to SE, and amount of KA received. (*p<0.05).

Robinson-Cooper, Davidson et al.
*Loss of presenilin 2 function age-dependently increases susceptibility to kainate-induced acute seizures and blunts hippocampal kainate-type glutamate receptor expression*
Supplemental Figures

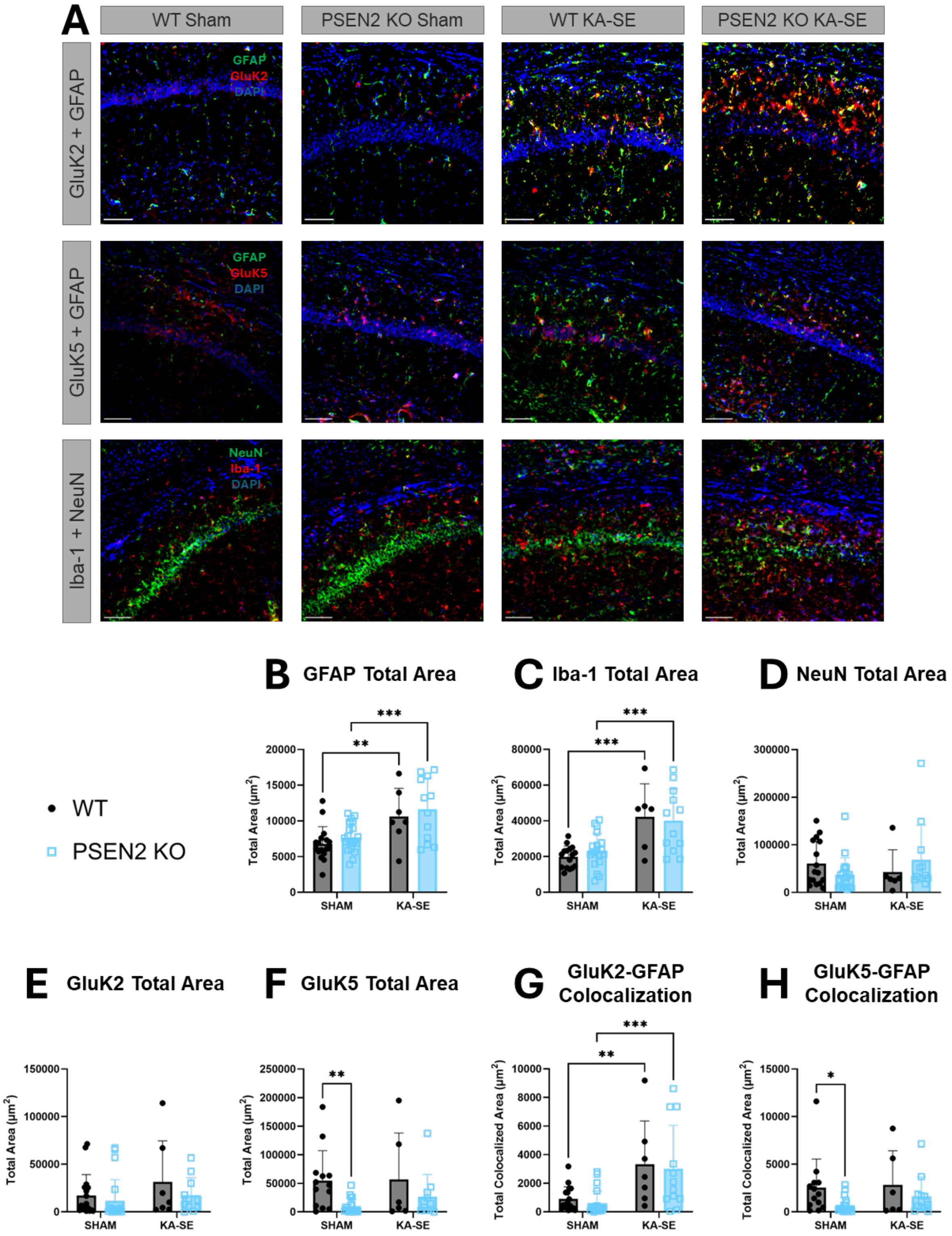

**Supplemental Figure 10 – Aged mouse histopathological changes in area CA1 of dorsal hippocampus**. Loss of normal PSEN2 function resulted in decreased bassline expression of GluK5 compared to age-matched WT mice aged 12-15 months. **A)** Representative images of GluK2 + GFAP, GluK5 + GFAP, and Iba-1 + NeuN staining at CA1 of WT and PSEN2 KO sham and KA-SE animals. **B)** GFAP expression in the CA1 region was increased in animals that received KA (Sham vs. KA-SE: $F (1, 53) = 20.6$, $p<0.0001$; WT Sham vs. KA-SE: **p = 0.006; PSEN2 KO Sham vs. KA-SE: ***p = 0.0006). **C)** Iba-1 expression in the CA1 region was increased in animals that received KA (Sham vs KA-SE: $F (1, 49) = 28.1$, $p<0.0001$; WT Sham vs. KA-SE: ***p = 0.0004; PSEN2 KO Sham vs. KA-SE: ***p = 0.0006). **D)** There was no difference in NeuN expression in the CA1 region between either treatment groups or genotypes. **E)** There was no difference in GluK2 expression in the CA1 region between either treatment groups or genotypes. **F)** Baseline GluK5 expression was significantly decreased in PSEN2 KO animals (WT vs. PSEN2 KO: $F (1, 45) = 7.69$, $p = 0.008$; Sham WT vs PSEN2 KO **p = 0.007) **G)** GFAP-GluK2 colocalization in the CA1 region was increased in animals that received KA (Sham vs. KA-SE: $F (1, 53) = 20.6$, $p<0.0001$; WT Sham vs. KA-SE: **p = 0.005; PSEN2 KO Sham vs KA-SE ***p = 0.0007). **H)** Baseline GFAP-GluK5 colocalization in the CA1 region is decreased in PSEN2 KO animals compared to WTs (WT vs. PSEN2 KO: $F (1, 46) = 4.97$, $p = 0.031$; Sham WT vs. PSEN2 KO *p = 0.026).

Robinson-Cooper, Davidson et al.
*Loss of presenilin 2 function age-dependently increases susceptibility to kainate-induced acute seizures and blunts hippocampal kainate-type glutamate receptor expression*
Supplemental Figures

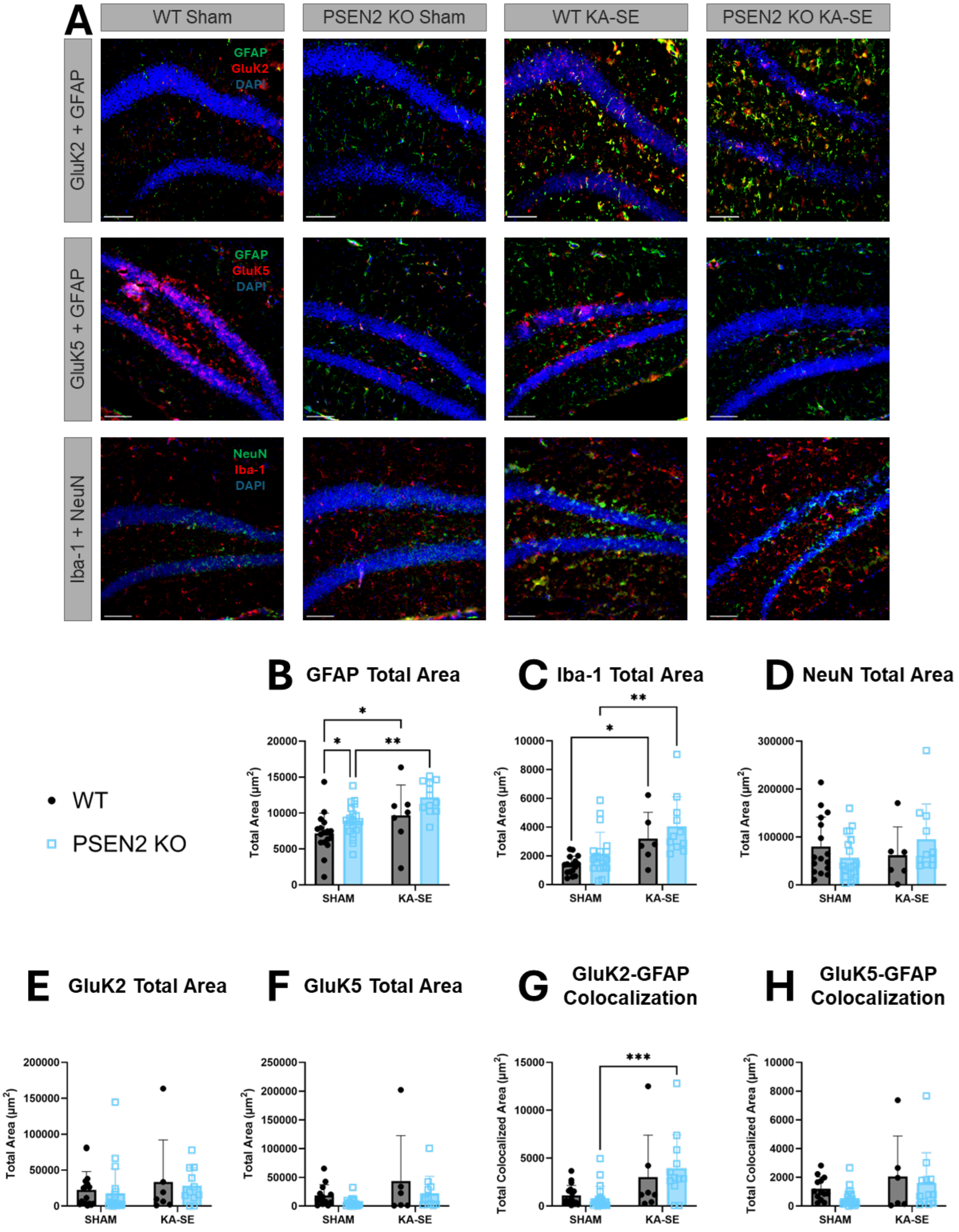

**Supplemental Figure 11. Aged mouse histopathological changes in dentate gyrus (DG) of dorsal hippocampus**. Loss of normal PSEN2 function resulted in decreased bassline expression of GluK5 compared to age-matched WT mice aged 12-15 months. A) Representative images of GluK2 + GFAP, GluK5 + GFAP, and Iba-1 + NeuN staining at DG of WT and PSEN2 KO sham and KA-SE animals. **B)** GFAP expression in the DG was increased in animals that received KA (Sham vs. KA-SE: F (1, 53) = 13.6, p = 0.005; WT Sham vs. KA-SE: *p = 0.044; PSEN2 KO Sham vs. KA-SE **p = 0.002) and in PSEN2 KO mice when compared to WTs (WT vs. PSEN2 KO: F (1, 53) = 7.62, p = 0.008; Sham WT vs. PSEN2 KO *p = 0.048) . **C)** Iba-1 expression in the DG was increased in animals that received KA (Sham vs KA-SE: F (1, 48) = 17.3, p = 0.0001; WT Sham vs. KA-SE: *p = 0.012; PSEN2 KO Sham vs. KA-SE **p = 0.001). **D)** There was no difference in NeuN expression in the DG between either treatment groups or genotypes. **E)** There was no difference in GluK2 expression in the DG between either treatment groups or genotypes. **F)** There was no difference in GluK5 expression in the DG between either treatment groups or genotypes. **G)** GFAP-GluK2 colocalization in the DG was increased in animals that received KA (Sham vs. KA-SE: F (1, 53) = 13.8, p = 0.005; PSEN2 KO Sham vs KA-SE ***p = 0.006). **H)** GFAP-GluK5 colocalization in the DG did not differ between either treatment groups or genotypes.

**A** 12–15-month-old r-value table

| | Acute Seizure Data | CA3 | | | | | CA1 | | | | | Dentate Gyrus | | | | |
|---|---|---|---|---|---|---|---|---|---|---|---|---|---|---|---|---|
| | | GFAP | GluK5 | GluK2 | NeuN | Iba-1 | GFAP | GluK5 | GluK2 | NeuN | Iba-1 | GFAP | GluK5 | GluK2 | NeuN | Iba-1 |
| WT | KA Received | 0.3 | 0.2 | 0.5 | 0.4 | -0.3 | 0.3 | 0.3 | **0.9** | 0.7 | -0.02 | 0.5 | 0.003 | 0.4 | 0.8 | -0.4 |
| WT | Time to 1st GTC | 0.6 | 0.3 | 0.7 | 0.3 | -0.2 | 0.5 | 0.3 | **0.9** | 0.6 | 0.05 | 0.7 | 0.08 | 0.6 | 0.7 | -0.4 |
| WT | Time to Status Epilepticus | 0.5 | 0.3 | 0.7 | 0.4 | -0.2 | 0.5 | 0.3 | **0.9** | 0.6 | 0.01 | 0.7 | 0.06 | 0.6 | 0.7 | -0.4 |
| WT | Seizure Burden | -0.04 | -0.5 | 0.3 | 0.6 | -0.4 | -0.07 | -0.4 | 0.3 | 0.2 | -0.3 | 0.06 | -0.5 | 0.3 | 0.5 | -0.2 |
| PSEN2 KO | KA Received | 0.1 | -0.1 | 0.1 | -0.006 | 0.006 | 0.1 | 0.2 | 0.1 | -0.06 | 0.2 | -0.008 | 0.1 | 0.1 | -0.05 | 0.03 |
| PSEN2 KO | Time to 1st GTC | 0.1 | 0.03 | 0.01 | -0.2 | -0.08 | 0.03 | 0.09 | 0.06 | -0.2 | 0.2 | 0.03 | 0.1 | -0.1 | -0.3 | 0.008 |
| PSEN2 KO | Time to Status Epilepticus | -0.01 | -0.04 | 0.08 | 0.06 | -0.02 | -0.08 | 0.1 | 0.05 | -0.007 | -0.07 | -0.1 | 0.1 | 0.006 | -0.05 | -0.07 |
| PSEN2 KO | Seizure Burden | **0.6** | -0.4 | 0.2 | 0.1 | 0.08 | **0.7** | -0.4 | 0.2 | 0.2 | 0.5 | 0.5 | -0.4 | 0.2 | 0.04 | 0.2 |

**B** GFAP Expression vs Acute Seizure Data Correlation

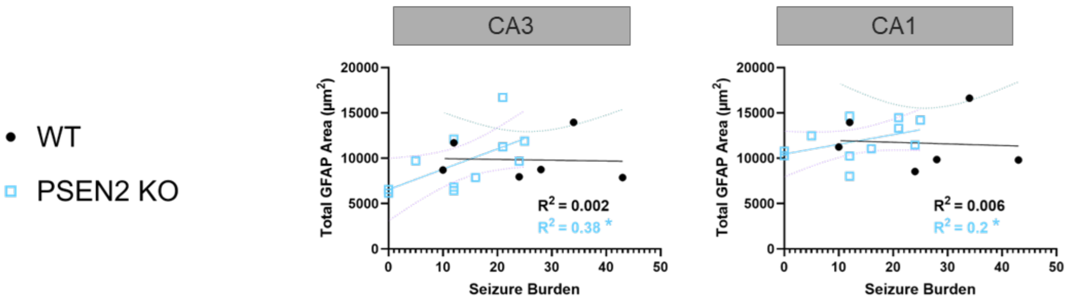


**C** GluK2 Expression vs Acute Seizure Data Correlation

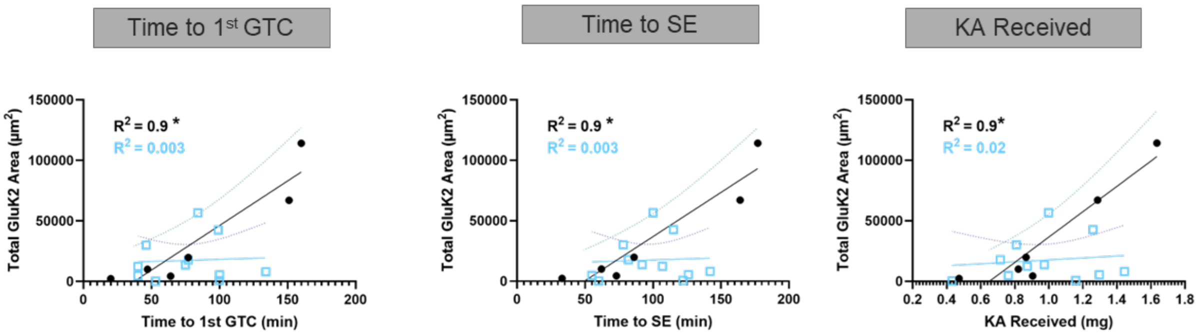


**Supplemental Figure 12. Acute seizure burden correlations to post-KA SE histology in aged mice.** Aged animal acute seizure data and histology correlation analyses. **A)** table of r-values obtained from each analysis. Red cells denote a significant correlation. **B)** Scatterplots of GFAP expression over seizure burden in CA3 and CA1. **C)** Scatterplots of total GluK2 area in the CA1 over time to 1st GTC, time to SE, and amount of KA received. (*p<0.05)
